# Cdc14 spatiotemporally regulates Rim4-mRNA complex assembly and stability during meiosis

**DOI:** 10.1101/2022.12.22.521673

**Authors:** Rudian Zhang, Wenzhi Feng, Suhong Qian, Shunjin Li, Fei Wang

## Abstract

In budding yeast, Rim4 sequesters a subset of meiotic transcripts and essentially suppresses their translation until being degraded at the end of meiosis I. We found that Rim4 loads mRNAs in the nucleus as a prerequisite for relocating Rim4 into the cytoplasm, where mRNAs protect Rim4 from autophagy. Nonetheless, the underlying mechanism remains unknown. Using genetic, biochemical, and cell imaging approaches, here, we revealed that phosphorylation states regulate Rim4’s intracellular interactions with mRNAs and the yeast 14-3-3 proteins Bmh1 and Bmh2. Our data showed that Rim4 forms a heterotrimeric complex with Bmh1 and Bmh2 via multiple phosphorylated sites with the consensus of a PKA kinase target. Remarkably, the Bmh1/2-Rim4 complex excludes mRNAs and is resistant to autophagy. We further found that Cdc14, a conserved cell cycle phosphatase, binds to a canonical Cdc14 docking site (PxL) in Rim4’s C-terminal low complexity domain (LCD) to de-phosphorylate Rim4 at multiple sites, resulting in Bmh1/2-Rim4 disassembly. Notably, before meiotic cell divisions, Cdc14 primarily resides in the nucleus, where mRNAs are transcribed. Therefore, Cdc14-triggered Bmh1/2 dissociation facilitates the nuclear Rim4 to target and sequester the nascent mRNAs. In contrast, during the meiotic divisions, Rim4-sequestered mRNAs are released for translation, while Cdc14 mediates Rim4-Bmh1/2 disassembly in the cytoplasm due to its temporary cytoplasmic relocation at the anaphases; subsequently, loss of protection from mRNAs and Bmh1/2 leads to autophagy-mediated Rim4 degradation at this stage. We conclude that phosphorylation states spatiotemporally regulate Rim4’s meiotic interactions, subcellular localization, and stability, regulated by Cdc14.

## Introduction

Meiosis aims to produce haploid gametes for sexual reproduction, featuring a single round of DNA replication followed by two sequential chromosome segregation, i.e., meiosis I and meiosis II. In *Saccharomyces cerevisiae* (budding yeast), meiosis is driven by a well-characterized transcriptional cascade, including starvation-induced *IME1* expression that supports the early meiotic transcription and *NDT80*, a mid-late meiotic transcription factor that controls about 300 genes (referred to as *NDT80* regulon) during meiotic divisions and the sporulation^1–6^. Regardless of being transcribed coordinately, the translation of these mRNAs is allocated to various stages of meiosis and sporulation as required by their distinct biological functions^7,8^. Accordingly, productive yeast gametogenesis is equipped with post-transcriptional regulation of the pre-existing mRNAs, primarily through RNA-binding proteins (RBP), which cooperatively engage both RNAs and proteins to exert various biological functions^9,10^.

In the past decade, technical advances have rapidly expanded the repertoire of RBPs in various organisms. A study of the *in vivo* mRNA interactomes in budding yeast captured 678 RBPs; many are involved in diverse biological processes^11^. However, molecular characterization and functional studies of the ribonucleoproteins (RNPs) that contain novel RNA-binding proteins remain technically challenging and slow-moving. Related to meiosis, several RBPs have been identified as meiosis-specific modulators in budding yeast, including *RIM4^12^, PES4*, and *MIP6*^7^, proposed to sequester a subset of meiotic transcripts to regulate their translation^7,12^. *PES4* and *MIP6* post-transcriptionally modulate a small portion of the ^~^300 genes induced by *NDT80 (NDT80* regulon) during meiotic prophase^7^. In contrast, *RIM4* is a general translation suppressor of the NDT80 regulon; the targets of Rim4 include main meiotic players such as *CLB3* and *AMA1*, a B-type cyclin for meiosis II and the activator of the meiotic anaphase-promoting complex (APC/C), respectively^12,13^. In line with Rim4 being a master meiotic translation suppressor, genetic deletion of *RIM4* abolished meiosis^14^.

Several studies proposed that Rim4 degradation is both required and sufficient to release Rim4-sequestered mRNAs for translation^13,15,16^. This scenario fits well with a ubiquitin-proteasome system (UPS)-mediated Rim4 degradation, which adapts a mechanism to remove protein molecules one by one, thereby will leave the mRNAs intact when Rim4 is degraded. However, we found that autophagy also actively degrades Rim4 during meiotic divisions and is required for *CLB3* translation^17^. Therefore, unless a mechanism releases mRNAs first, Rim4 degradation by autophagy, featuring autophagosome-mediated engulfment, would not spare Rim4-sequestered mRNAs. Indeed, we found that the Rim4-mRNA Ribonucleoprotein (RNP) complex resists autophagy, and mRNAs dissociation is a prerequisite of Rim4 degradation by autophagy, thereby sparing the mRNAs for translation (accompanying manuscript). However, it remains unknown how the assembly and disassembly of Rim4-mRNA RNP are programmed by meiosis.

Almost every facet of RNA biology, e.g., RNA transport, degradation, and translation, is regulated by phosphorylation. Along meiosis progression, a programmed oscillation of kinase and phosphatase activities, e.g., by CDK1(Cdc28) kinase and the Cdc14 phosphatase, primarily coordinates the timing of meiotic events. Therefore, a temporal switch on Rim4-mRNA interaction during meiosis could base on phosphorylation. As one clue, before its rapid degradation, Rim4 is cumulatively phosphorylated during meiosis by Ime2, a CDK2-like kinase^12^. Interestingly, a hypo-phosphorylated Rim4 mutant (Rim4[47A]) exhibited enhanced self-assembly and binding to *CLB3* mRNA, a model Rim4 substrate^15^. This finding implies that phosphorylation states affect Rim4-Rim4 and Rim4-mRNAs interactions. Of course, it is unreliable to conclude based on a hypo-phosphorylation mimic, which contains substantial replacement of 47 S/T residues by alanine (Rim4[47A]). Thus, the physiological nature of Rim4 phosphorylation and its meiotic modulators remain unclear.

Here, we elucidated how phosphorylation regulates Rim4-mRNA assembly and disassembly in the context of timed Rim4 degradation by autophagy during meiotic divisions. Specifically, we defined multiple primary phosphorylation sites (P-site) on Rim4 that are physiological targets of PKA (cAMP-dependent protein kinase) and Cdc14, a cell cycle phosphatase. Our data demonstrate that Cdc14 de-phosphorylates Rim4-p and triggers Rim4 dissociation from Bmh1/2, the yeast 14-3-3 proteins. This reaction leads to 1) autophagy-resistant Rim4-mRNA complex assembly when Cdc14 resides in the nucleus and 2) autophagic Rim4 degradation when Cdc14 activity transiently relocates to the cytoplasm during meiotic divisions.

## RESULTS

### Phosphorylation regulates the subcellular distribution of Rim4

We demonstrated in our accompanying manuscript that Rim4 enters the nucleus to pick up nascent mRNAs, an event required for Rim4 nuclear export. To examine whether phosphorylation regulates Rim4-mRNA interaction, we searched for P-sites on Rim4 that influence its subcellular distribution. To this end, we evenly divided the Rim4 sequence (713 residues) into nine regions (R1 to R9); next, all Serine (S) and Threonine (T) residues in each region were mutated to Glutamic acid (E) to mimic a hyper-phosphorylation state, or to Alanine (A) to mimic a hypo-phosphorylation state (Fig. 1A, top). As proof of interfered Rim4-mRNA interaction, the E or A mutations in R2, R3, and R5, which contain an RNA-binding RRM domain, block meiotic DNA replication (Fig. 1A) and sporulation (Fig. 1B). In contrast, meiotic DNA synthesis and sporulation tolerate E/A mutations in R1, R4, R8, and R9, an indication of lacking major phosphorylation-regulated Rim4 function in these regions (Fig. 1A; 1B). Interestingly, the sporulation of cells expressing R6-E and R7-A appeared to be the lowest except for the RRM-containing mutants (Fig. 1B), implicating that phosphorylation states of S/T residues in these regions physiologically affect Rim4.

**Figure 1.**
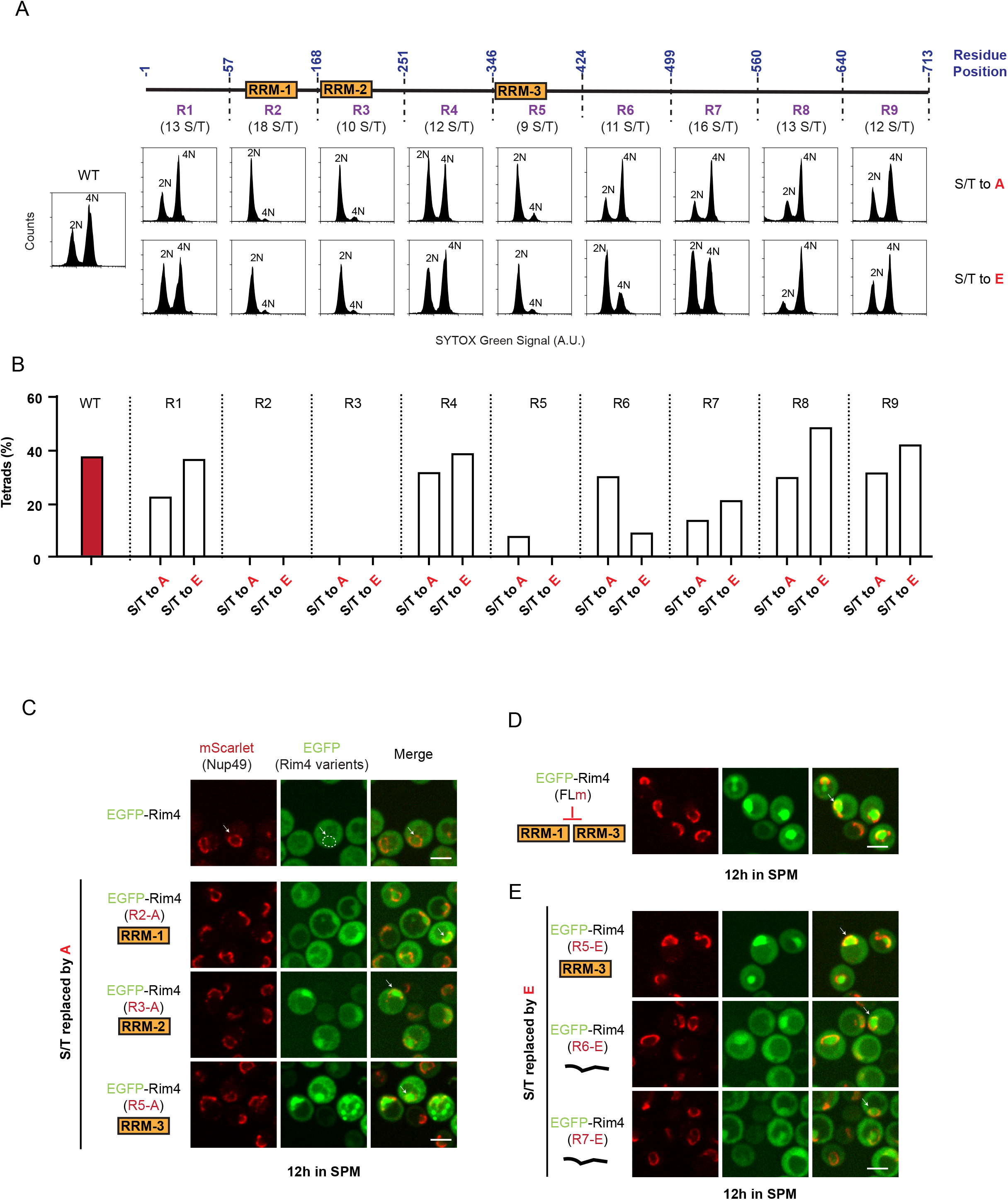
Phosphorylation regulates the subcellular distribution of Rim4. A) <u>Top</u>, the schematic of systematic Rim4 mutagenesis. In Each indicated region, all Serine (S) and Threonine (T) residues were replaced by Alanine (A) or Glutamic Acid (E). The yellow boxes indicate RRM-1, RRM-2, and RRM-3. <u>Bottom</u>,flow cytometry (FACS) analysis of the DNA content (2N versus 4N) in the indicated Rim4 variants; cells were collected for measurement at 12 h in SPM before *NDT80* induction (*GAL-NDT80* system). B) The sporulation efficiency of indicated Rim4 variants, presented as the tetrads percentage; more than 300 cells were counted for each strain. C-E) The presentative FM images showing the intracellular distribution of Rim4 variants with EGFP tagged at the N-terminus (green); Nup49-mScarlet marks the nuclear membrane (red). The arrow (white) indicates the nucleus; the dashed line marks a nuclear membrane’s outline. Data are from three independent experiments. The scale bar is 5 μm.

Next, by fluorescence microscopy (FM) analysis of cells arrested at prophase I due to lack of *NDT80* induction (GAL-NDT80–synchronized meiosis)^18^ (Fig. S1A), we revealed that RRM-containing R2-A, R3-A, R5-A, and R5-E caused spatial Rim4 (*pRIM4:EGFP-Rim4[variants]*)accumulation to various degrees in a subcellular compartment marked as the nucleus by Nup49 (*pNUP49:Nup49-mScarlet*), a nuclear pore complex component (Fig. 1C; S1B). The behavior of these mutants is reminiscent of the mRNA-binding defective Rim4 mutant (GFP-Rim4[FLm], i.e., the F_139_L-F_349_L mutant, in which we mutated two mRNA binding sites in the conserved RRM1 and RRM3^19–21^ (Fig. 1D). This result further supports that mRNA binding is required for efficient Rim4 nuclear export. Interestingly, R2-E and R3-E did not cause Rim4 nuclear retention as R2-A and R3-A did, suggesting that phosphorylation states of the RRM1 and RRM2 and their flanking sequences might directly modulate Rim4-mRNA interactions. Nevertheless, the substantial residue replacement in the RRM-containing region, e.g., 18 out of 111 residues changed in R2 (RRM1), might introduce phosphorylation-independent effects that artificially alter the conserved RRM domain structure. Besides, some S/T residues in a folded RRM might not even be accessible to a kinase or phosphatase. Due to these concerns, we avoided mutating the RRMs in the following experiments unless mutagenesis of a specific residue was pursued.

Unlike an RRM, the C-terminal Rim4 low complexity domain (LCD) that contains the R6-R9 region is intrinsically disordered and highly accessible to enzyme activities (Fig. S1C)^22^; and previous proteomic analysis has identified cumulative phosphorylation in the LCD region along meiosis^15^. Intriguingly, among the examined mutants, R6-E and R7-E also caused Rim4’s nuclear retention (Fig. S1B; 1E), correlated to their reduced DNA replication and sporulation efficiency (Fig. 1A; 1B). In contrast, the hypo-phosphorylation-mimicking R6-A and R7-A did not cause Rim4’s nuclear retention (Fig. S1B). These findings collectively suggest that the LCD domain, albeit lacking structured RRM, regulates Rim4-mRNA interaction. In particular, dephosphorylation of a specific set of S/T residues in the R6 and R7 regions encourages Rim4-mRNA complex assembly in the nucleus.

### Bmh1 and Bmh2 chaperones phosphorylated Rim4 in a heterotrimeric complex

To probe how phosphorylation regulates Rim4-mRNA, next, we affinity-purified Rim4-FLAG (IP: α-FLAG) from the lysates derived from meiotic cells (12h in SPM) before cellular Rim4 removal occurred (Fig. 2A) and analyzed the Rim4 interactome by Mass Spectrometry (Fig. 2B). The Bmh1, Bmh2, and Pab1 are top interactors with Rim4 in both cytosolic fraction (S_100_: 100,000 × g, 49 min) and a detergent-solubilized rough membrane fraction (P_100_) that contains nucleus and aggregates. (Fig. 2A; 2B). Strikingly, in the cytosolic fraction cleared by ultracentrifugation (S_100_), Rim4-FLAG forms a stable heterotrimeric complex with Bmh1 and Bmh2 with a 1:1:1 stoichiometry, as determined by Mass Photometry (MP), which characterizes the mass distribution of the three proteins (Fig. 2C). Moreover, based on MP, the Rim4-Bmh1/2 complex is free of any other macromolecular mass, e.g., Pab1 or mRNAs. Thus, intracellular Rim4 forms multiple complexes; one exclusively contains a Bmh1/2 heterodimer that chaperones a Rim4 monomer.

**Figure 2.**
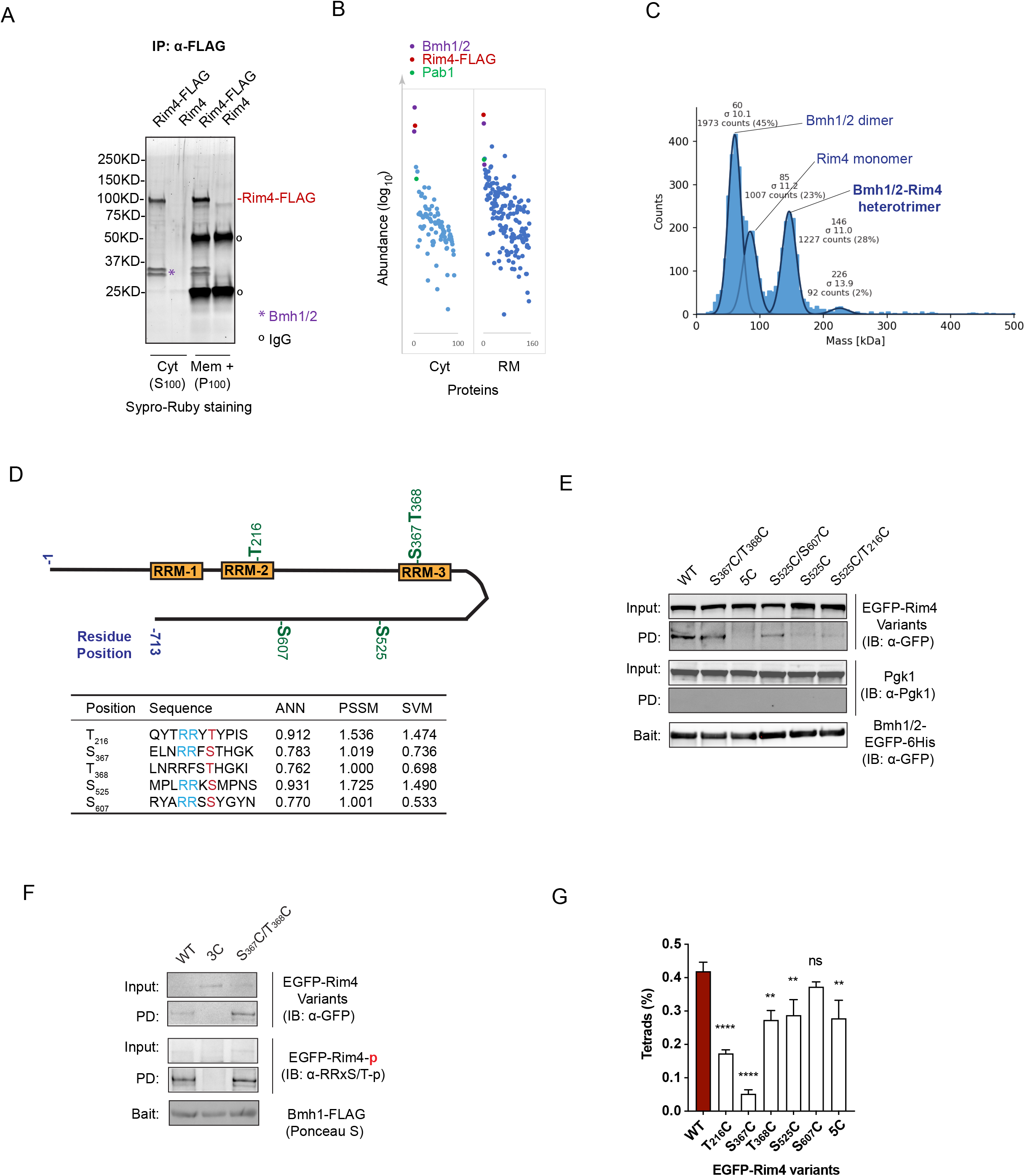
14-3-3 proteins Bmh1 and Bmh2 chaperones phosphorylated Rim4 in a heterotrimeric complex. A and B) Identification of Rim4 interactome by IP and mass spectrometry A) IP of Rim4-FLAG in the lysates of cells collected at 12 h in SPM. The immunoprecipitated Rim4-FLAG and its interactions were resolved by SDS-PAGE and stained with SYPRO-Ruby. The purple asterisk marks the Bmh1 and Bmh2 doublet, with Bmh1 at the bottom; black circles indicate the IgG. B) Graph of the abundance of Rim4 interacting proteins identified by Mass Spectrometry. Rim4, Bmh1, Bmh2, and Pab1 are depicted in color among the most abundant proteins in the IP-purified complex of Rim4-FLAG, as shown in (A). C) Mass Photometry analysis of cytosolic Rim4-FLAG complex achieved in (A). The histogram (light blue) shows the number of contrast events related to a given molecular mass. The solid lines are Gaussian fits to the peaks. The mean and standard deviation (σ) of each Gaussian curve is shown above it, as is the number of counts accounted for the peak. D) Schematic of 14-3-3 protein (Bmh1/2) binding sites (top) on Rim4 predicted by the 14-3-3-Pred program^26^ (bottom). The prediction was made with three methods: Artificial Neural Network (ANN, cut-off = 0.55), Position-Specific Scoring Matrix (PSSM, cut-off = 0.8), and Support Vector Machine (SVM, cut-off = 0.25). The listed sites pass all three methods. Phosphorylation of the marked S/T residue (red) is required for Bmh1/2 binding to the site. The RRxS/T sequence is the consensus of PKA targets. E) The Ni-NTA beads immobilized Bmh1/2-EGFP-6×His proteins pull down EGFP-tagged Rim4 variants, i.e., WT, single (S_525_C), double (S_367_C/T_368_C, S_525_C/S_607_C, S_525_C/T_216_C) or quintuple (5C) mutants, from the prophase I cell lysate. The elutes with Imidazole are analyzed by immunoblotting with indicated antibodies. Pgk1 serves as a negative control. F) Co-immunoprecipitation of indicated Rim4 variants, i.e., WT, double (S_367_C/T_368_C) or triple (T_216_C/S_525_C/S_607_C = 3C) mutants, with Bmh1-FLAG (p*BMH1: Bmh1-FLAG*) in prophase I cell lysates. The elutes with 3xFLAG peptides are analyzed by immunoblotting, with α-GFP antibody to detect EGFP-Rim4 variants or α-RRxS/T-p antibody to detect phosphorylated EGFP-Rim4 variants. G) The sporulation efficiency of indicated Rim4 variants, presented as the tetrad percentage; WT cells are compared with single (T_216_C, S_367_C, T_368_C, S_525_C, S_607_C) or quintuple (5C) mutant Rim4. ****: p<0.0001, **: 0.001<p<0.01, ns: no significance. Data are from three independent experiments; more than 300 cells are counted for each.

To our delight, Bmh1 and Bmh2 are the yeast 14-3-3 proteins and predominantly bind consensus that contains a central phosphorylated Serine (S-p) or Threonine (T-p)^23,24^. Typically, de-phosphorylation at that single S-p/T-p is sufficient to dissociate 14-3-3 proteins from their target^25^. With each monomer consisting of nine α-helices to form a highly conserved amphipathic groove as the phosphorylation-binding pocket, the heterodimer and homodimer of Bmh1 and Bmh2 often simultaneously accommodate two peptides in its binding groove to achieve high affinity^25^. Rim4 contains four highly scored putative Bmh binding sites (BBS) with an RRxS/T consensus: one in RRM2 (T_216_-p), one in RRM3 (S_367_-p/T_368_-p), and two in the LCD (S_525_-p in R-7; S_607_-p in R-8), predicted by 14-3-3-Pred (Fig. 2D)^26^. All four putative BBSs are likely accessible to Bmh binding based on protein structure prediction; three BBSs are unstructured (S_525_-p; S_607_-p) or in a loop (T_216_-p), while the S_367_/T_368_ site is exposed to the surface of folded RRM3 structure (Fig. S2A). Therefore, the stable 1:1:1 Rim4-Bmh1-Bmh2 complex indicates that more than two of the predicted BBSs are valid, and multiple states of the Rim4-Bmh1/2 complex are possible (Fig. S2B).

Next, we used site-specific mutagenesis to substitute the central S/T residues of the four BBSs with Cysteine (C). Due to the structural similarity, the replacement of S by C is neutral while lacking phosphorylation ability. By a pull-down with bead-immobilized recombinant Bmh1/2 (BMH1/2-GFP-6×His), we found that the S_525_C, S_525_C/T_216_C, and S_525_C/S_607_C mutants reduced, while mutation of all five S/T residues (Rim4[5C]) abolished Rim4-Bmh1/2 interaction in the prophase I cell lysate (Fig. 2E). In addition, the T_216_C/S_525_C/S_607_C triple mutant (Rim4[3C]) almost entirely abolished endogenous Rim4-Bmh interaction (Fig. 2F), supporting that at least two BBSs are required for stable Rim4-Bmh1/2 interaction. Furthermore, phosphorylation of Rim4 at RRxS/T (BBS) sites in prophase I cell lysates, confirmed by immunoblotting (IB: *a-*RRxS/T-p), almost vanished in the triple mutant Rim4(3C) but not in Rim4(S_367_C/T_368_C) (Fig. 4F), indicating that T_216_, S_525_, and S_607_ are minimally involved in Bmh1/2-Rim4 interaction.

In cells at prophase I, mutation of the S_367_/T_368_ site (Rim4[S_367_C/T_368_C]) did not significantly affect Rim4-Bmh interaction, like Rim4(S_525_C) did (Fig. 2E). Therefore, we proposed that S_525_C is critical for Rim4-Bmh1/2 interaction, presumably in multiple states (Fig. S2B). Nonetheless, S_367_C and T_368_C exhibited reduced sporulation efficiency (Fig. 2G), as T_216_C and S_525_C did, indicating that phosphorylation at the S_367_/T_368_ site might occur at a different meiotic stage or only transiently facilitate the transition between different heterotrimeric Bmh-Rim4 states (Fig. S2B). We speculate that Bmh1/2 might switch between BBSs to regulate different aspects of Rim4 activity, guided by the phosphorylation states of S/T residues at each BBS (Fig. S2B). As support, sporulation of Rim4(5C) cells is better than most single BBS mutations, e.g., T_216_C and S_367_C (Fig. 2G), suggesting that effects of Bmh binding on Rim4 at different BBSs can partially cancel each other. We will discuss this scenario later.

### PKA-stimulated Rim4-Bmh1/2 interaction antagonizes Rim4-mRNA assembly

Using Mass Photometry analysis, we defined a cytosolic Rim4-Bmh1-Bmh2 heterotrimer, lacking apparent RNAs (Fig. 2C). In cells at prophase I, we found that Rim4(5C) (pRIM4:GFP-*Rim4[5C]*), the mutant lacking Bmh1/2 binding, resides in the nucleus less than the wild type Rim4 (Fig. 3A), an indication of faster Rim4-mRNA complex formation and nuclear export in the absence of Bmh1/2. These findings, in line with BBSs overlapping with the RRMs on Rim4 (Fig. 2D), suggest that Bmh1/2 senses phosphorylation at the BBSs to restrict Rim4-mRNA complex formation.

**Figure 3.**
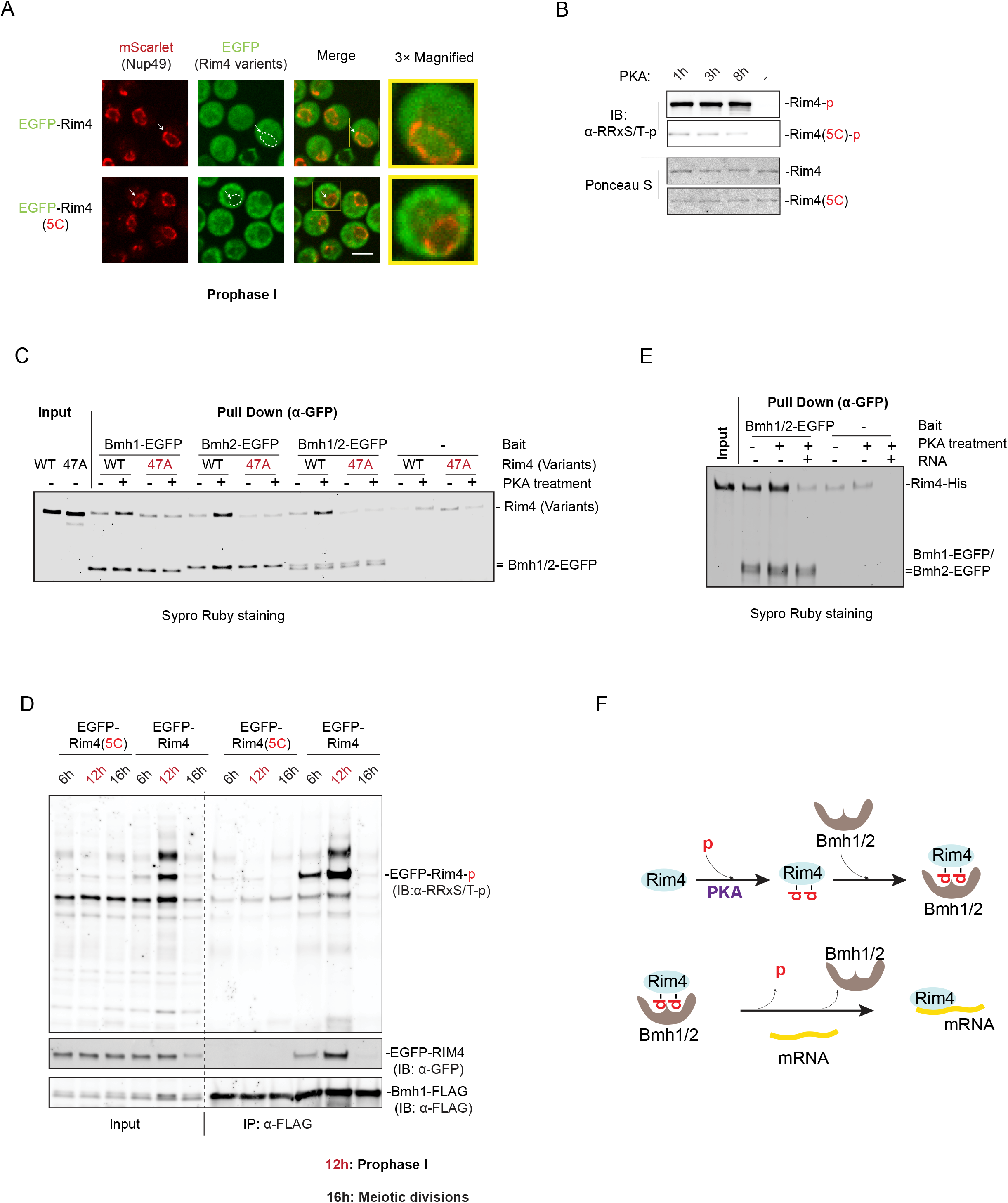
PKA-stimulated Rim4-Bmh1/2 interaction antagonizes Rim4-mRNA assembly. A) The presentative FM images showing the intracellular distribution of Rim4 variants with EGFP tagged at N-terminus (green); Nup49-mScarlet marks the nuclear membrane (red). The 3X zoom-in images (right) show the nuclear signal of WT and Rim4(5C). The arrow (white) indicates the nucleus; the dashed line marks the outline of a nuclear membrane. Data are from three independent experiments. The scale bar is 5 μm. B) PKA catalytic subunit (Sigma, P2645) mediated Rim4 phosphorylation at the RRxS/T sites was conducted *in vitro*, using recombinant Rim4-6×His; the recombinant Rim4(5C)-6×His acted as a negative control. Shown are IB analyses of the proteins after PKA treatment for indicated time at room temperature, using an α-RRx-S/T-p antibody that specifically recognizes phosphorylated RRx-S/T sites. Ponceau S staining shows protein levels of Rim4 variants in the reaction. C) After one hour of PKA or mock treatment, the indicated Rim4 variants were pulled down by beads-immobilized Bmh1/2-EGFP-6×His (α-EGFP). Rim4(47A) is a mutant with the 47 S/T residues in the C-terminal LCD of Rim4 simultaneously mutated to A. The samples were resolved on SDS-PAGE and visualized after Sypro Ruby staining. D) IB analysis of EGFP-Rim4, or EGFP-Rim4(5C), phosphorylation at BBSs and its coimmunoprecipitation with Bmh1-FLAG (IP: α-FLAG) in cells collected from the early meiotic stage (6h in SPM), prophase I (12h in SPM, without NDT80 induction), and meiotic divisions (16h in SPM, four hours after induced NDT80 expression). Phosphorylation at BBSs was detected by an α-RRxS/T-p antibody; the EGFP-tagged Rim4 variants were detected by an α-GFP antibody. Note that Rim4 phosphorylation and its interaction with Bmh1 reached a peak at prophase I. E) The total yeast RNAs inhibit PKA-stimulated Rim4-Bmh1/2 interaction. Applied RNAs, at the concentration of 50 ng/ μL, were pre-mixed with the PKA-treated Rim4 protein before incubation with Bmh1/2. Sypro Ruby staining of the input and pull-down samples resolved by SDS-PAGE are shown. F) A model showing phosphorylation (by PKA) state at the Rim4 BBSs determines Rim4 interaction. Phosphorylated Rim4 forms a complex with Bmh1/2; de-phosphorylated Rim4 binds mRNAs.

**Figure 4.**
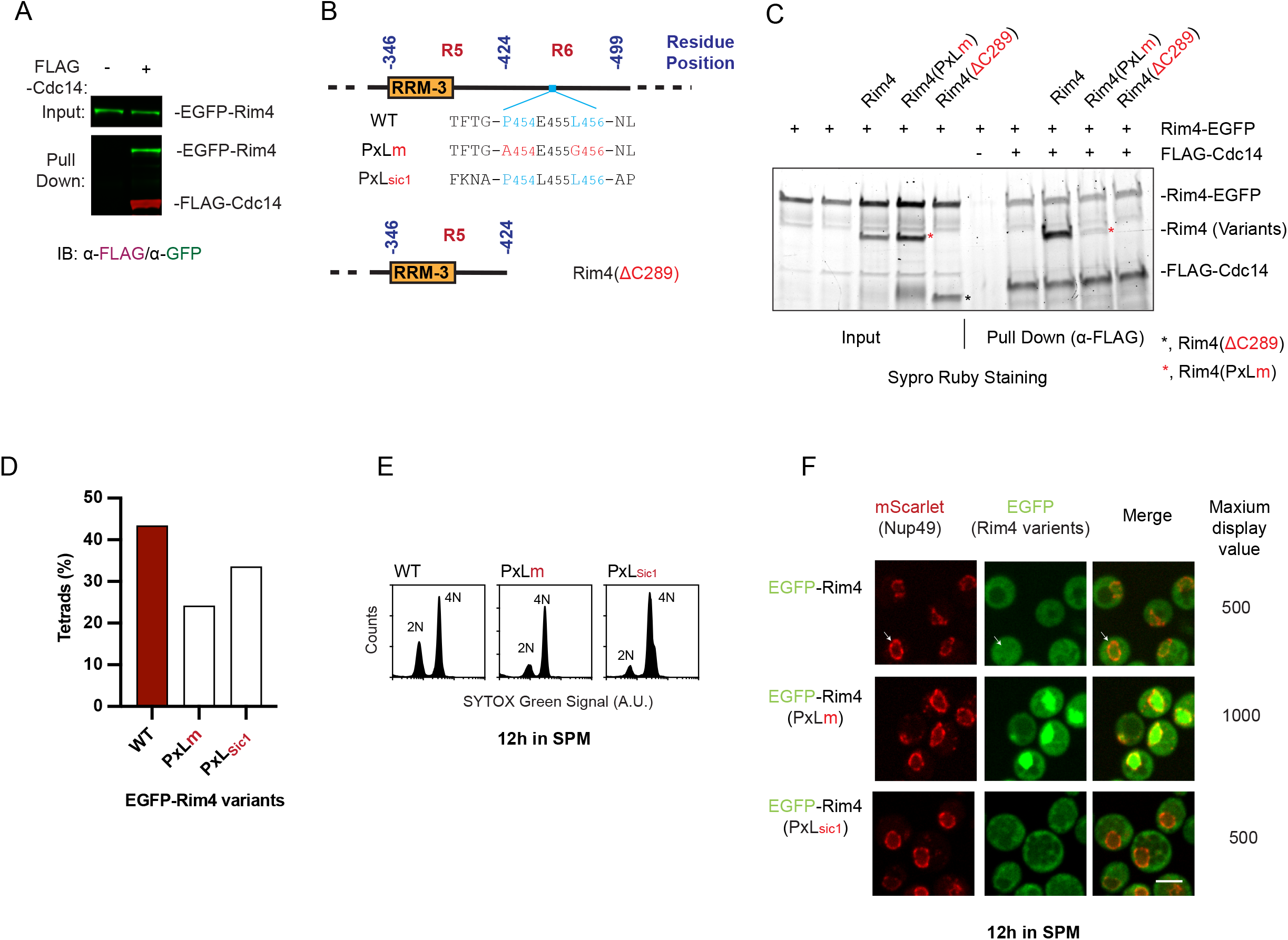
Cdc14 binds Rim4 to assist its nuclear export. A) Bead-immobilized Cdc14-FLAG (α-FLAG) pulls down EGFP-Rim4 (pRIM4: *EGFP-Rim4*) from prophase I cell lysates controlled by empty α-FLAG beads. B) Top, schematics of Rim4, harboring a wild type (WT), or modified Cdc14 docking site (PxLm and PxL_sic1_). Bottom, schematics of Rim4 with the C-terminal 289 residues truncated (Rim4 [ΔC289]). C) Bead-immobilized Recombinant Cdc14-FLAG (α-FLAG) pulls down recombinant Rim4 variants, including Rim4-EGFP and Rim4-6×His. The Rim4(PxLm)-6×His and Rim4(ΔC289)-6×His binds poorly with Cdc14-FLAG in the presence of Rim4-EGFP (competitor). Red asterisk: Rim4(PxLm)-6×His, black asterisk: Rim4(ΔC289)-6×His. D) The sporulation efficiency of indicated Rim4 variants, presented as the tetrads percentage; more than 300 cells were counted for each strain. E) The DNA content in cells at 12h in SPM, measured by FACS; the strains express either Rim4, Rim4(PxLm), or Rim4(PxL_sic1_). F) The presentative FM images showing the intracellular distribution of Rim4 variants with EGFP tagged at N-terminus (green), including Rim4, Rim4(PxLm), and Rim4(PxL_sic1_); Nup49-mScarlet marks the nuclear membrane (red). The maximum display value (EGFP) was set to 1000 for Rim4(PxLm) and 500 for Rim4 and Rim4(PxL_sic1_). The arrow (white) indicates the nucleus; the dashed line marks the outline of a nuclear membrane. Data are from three independent experiments. The scale bar is 5 μm.

Next, we decided to examine this mechanism with purified components *in vitro*. Our primary goal was to reconstitute 1) the phosphorylation at the BBSs of Rim4 (Rim4_BBS_-p), 2) the formation of Bmh1/2-Rim4_BBS_-p complex, and 3) a possible competition between mRNAs and Bmh1/2 on Rim4 binding. First, we noticed that the S/T residues at the Rim4 BBSs are all canonical PKA kinase targets, with a consensus of RRxS/T-p^27,28^. Accordingly, using a PKA substrate antibody that only recognizes phosphorylated RRxS/T (IB:α-RRxS/T-p), we showed that PKA remarkably phosphorylates recombinant Rim4, but not Rim4(5C) at the BBSs (Fig. 3B) *in vitro*. Next, we showed that resin-immobilized GFP-tagged Bmh1, Bmh2, or Bmh1/2 (IP: α-GFP) pulled down Rim4_BBS_-p after PKA treatment (Fig. 3C). In contrast, Rim4 failed to interact with Bmh proteins above the noise level (Fig. 3C). Moreover, we found that Rim4(47A), in which Alanine (A) replaces 47 S/T residues including S_525_ and S_607_ in the LCD region, did not engage Bmh1/2 even after PKA treatment (Fig. 3C). These results demonstrate that simultaneous phosphorylation at the Rim4 BBSs that includes S_525_ is both sufficient and required for stable Bmh-Rim4 interaction.

Previous studies found that Ime2 phosphorylates Rim4 during meiosis, although the specific functional P-sites of Ime2 activity were unidentified^12^. Unlike PKA, Ime2 treatment did not efficiently phosphorylate recombinant Rim4 *in vitro* (Fig. S3A). *In vivo*, the cellular Rim4_BBS_-p level and Rim4-Bmh interaction reach the peak in the wild type but not in the Rim4(5C) cells at prophase I as determined by IB analysis using PKA substrate antibody (IB:α-RRxS/T-p), following IP of Bmh1-FLAG (IP: α-FLAG) (Fig. 3D; S3B). Based on the literature, Ime2-mediated Rim4 phosphorylation mainly accumulates during the meiotic divisions^12,29^ when the cellular Rim4_BBS_-p level declines (Fig. 3D). Therefore, our data suggest that PKA primarily mediates Rim4 phosphorylation at the BBSs to regulate Rim4-Bmh1/2 interaction (Fig. 3F, top).

Now that we prepared pure Rim4_BBS_-p *in vitro* that can robustly bind Bmh proteins, we examined whether mRNA and Bmh proteins could simultaneously bind Rim4_BBS_-p. Remarkably, the total yeast RNAs at a concentration of 50 ng/μl, premixed with Rim4 _BBS_-p, effectively expel Bmh1/2 from Rim4 _BBS_-p (Fig. 3E), indicating that mRNAs compete with Bmh1/2 on Rim4 docking sites. Notably, Bmh1 and Bmh2 are among the most abundant yeast proteins. Therefore, *in vivo*, the cellular mRNAs on their own are unlikely to release Bmh1/2 from phosphorylated Rim4, consistent with our finding that Rim4 forms stable heterotrimeric complexes with Bmh1/2 (Fig. 2A). Thus, we hypothesize that de-phosphorylation at the BBS sites dissociates Bmh1/2 from Rim4, as a prerequisite of efficient Rim4-mRNA complex assembly in the cell (Fig. 3F, bottom).

### Cdc14 binds Rim4 to assist its nuclear export

Our model predicts that a phosphatase, ideally in the nucleus, is required for efficient Bmh1/2 dissociation and the formation of Rim4-mRNA RNP before its nuclear export. So, we examined Cdc14, which primarily resides in the nucleus except for the anaphases. Encouragingly, recombinant Cdc14 (Cdc14-FLAG) pulls down Rim4 (IP: α-FLAG) in the prophase I cell lysates (Fig. 4A). Rim4 harbors one putative Cdc14 docking PxL site, P_454_E_455_L_456_, in the R6 region (Fig. 4B)^30^. Next, we confirmed that Rim4-Cdc14 interaction is via the P_454_E_455_L_456_ site. Bead-immobilized (IP: α-FLAG) recombinant Cdc14 (Cdc14-FLAG) pulled down recombinant Rim4 and Rim4-GFP, but not Rim4(ΔC289) or Rim4(PxLm), in which the P_454_E_455_L_456_ site was removed by truncation of 289 residues from C-terminus (ΔC289) or by point mutations at the A_454_E_455_G_456_ residues (PxLm) (Fig. 4B; 4C).

*In vivo*, the Rim4(PxLm) impaired sporulation (Fig. 4D), albeit supporting normal meiotic DNA replication (Fig. 4E). The reduced sporulation can be partially restored by replacing the Rim4 PxL site and its flanking residues (TFTGPELNL) with a Cdc14 docking site derived from Sic1 (FKNAPLLAP)^30^ (Fig. 4B, Rim4[PxL_sic1_]; 4D). Strikingly, by FM analysis of cells at prophase I, we found that the presence of Rim4(PxLm) (*pRIM4:GFP-Rim4[PxLm]*) mutant in the nucleus is greatly enriched, while Rim4(PxL_Sic1_) (*pRIM4:Rim4[PxL_Sic1_]*) supports WT-like subcellular Rim4 distributes (Fig. 4F). We noticed that the texture of cytoplasmic Rim4(PxL _Sic1_) under FM is somewhat decorated with more puncta than that of the WT Rim4 (Fig. 4F), reminiscent of some hypo-phosphorylation mutant of Rim4, e.g., R7-A (Fig. S1B), suggesting that Sic1 PxL might cause aberrant Rim4 de-phosphorylation by Cdc14 due to changed Cdc14-Rim4 interaction. Accordingly, Rim4 with Sic1 PxL site is less functional (Fig. 4D). Remarkably, Rim4(PxLm) exhibited nuclear retention phenotype as early as at the meiosis entry (Fig. S4A), indicating that Cdc14 assists Rim4 nuclear export upon Rim4’s meiotic translation, via the P_454_E_455_L_456_ site.

### Cdc14 de-phosphorylates Rim4 to release Bmh1/2

The nuclear retention phenotype of Rim4(PxLm) suggests that Cdc14 stimulates Rim4-mRNA complex assembly in the nucleus. So, we hypothesized that the S/T residues at BBSs are Cdc14 targets and that Cdc14 triggers Rim4-Bmh1/2 disassembly. To test this hypothesis, we supplemented 0.1 μg/μL Cdc14 to the prophase I cell lysate, in which we observed a high level of Rim4_BBS_-P and strong Rim4-Bmh1/2 interaction (Fig. 3D). After incubation for 20 min at room temperature, Bmh1/2 pulls down less Rim4 from Cdc14-treated cell lysates, as compared to the mock treatment (Fig. 5A; IB: α-GFP); moreover, the phosphorylation at the BBSs of pull-down Rim4 disappeared after Cdc14 treatment as determined by IB (α-RRxS/T-p) (Fig. 5A). Next, in the Rim4(PxLm) mutant cells arrested at prophase I, we found that the level of phosphorylation at BBSs is greatly enhanced (Fig. 5B). These results indicate that Cdc14 de-phosphorylates the BBSs on Rim4, leading to Bmh1/2 release.

**Figure 5.**
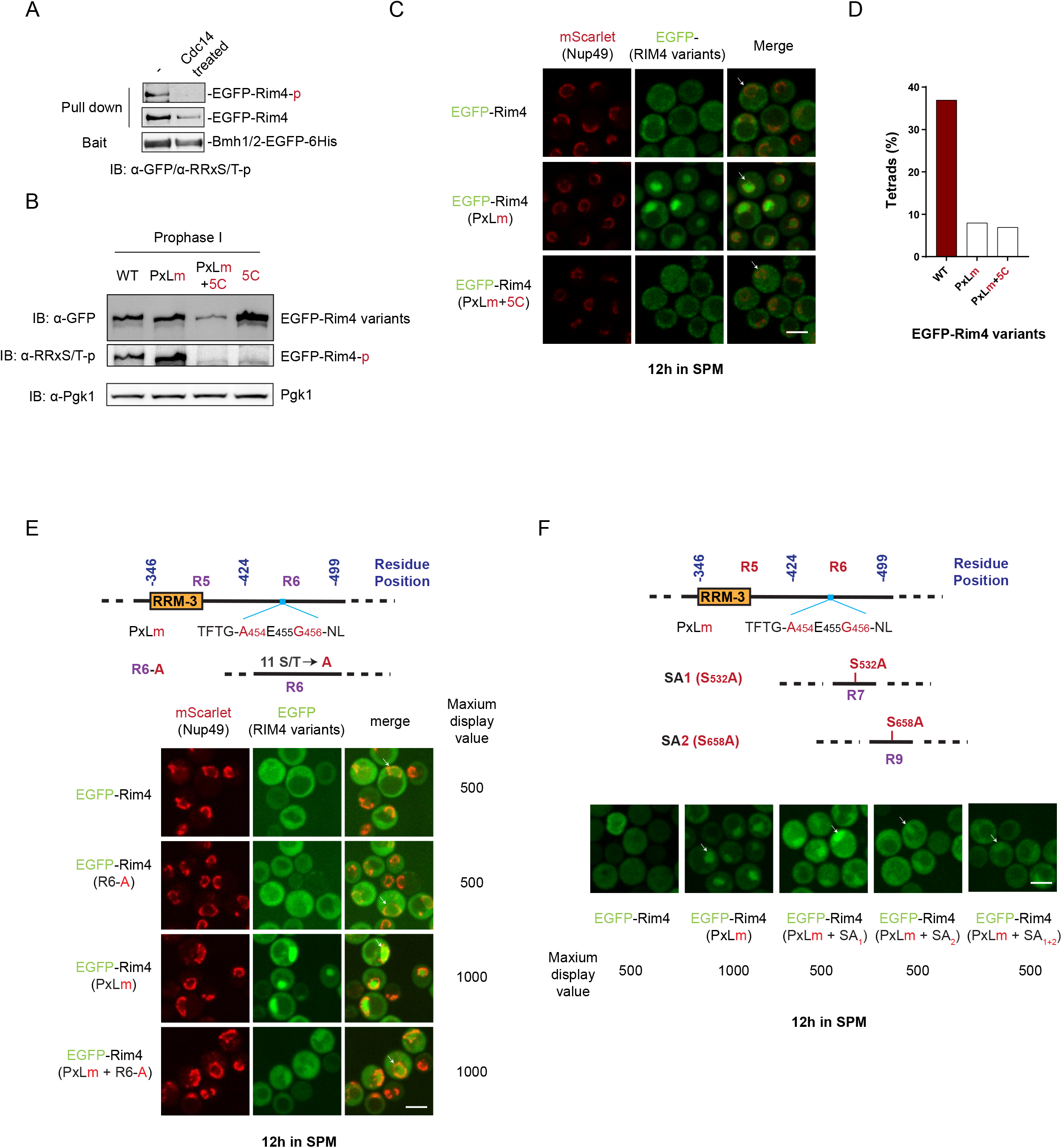
Cdc14 de-phosphorylates Rim4 to regulate its intracellular interaction and subcellular distribution. A) IB analysis of NTA-resin-immobilized recombinant Bmh1/2-EGFP-6×His pulling down endogenous EGFP-Rim4 from prophase I cell lysates. Before incubation with Bmh1/2-EGFP-6×His, the cell lysates were treated with Cdc14 (1.5 μM) for 30 minutes at room temperature or mock treatment. B) IB analysis of prophase I (12h in SPM) cell lysates derived from indicated Rim4 variants (EGFP-tagged), with indicated antibodies. Note that phosphorylation of Rim4(PxLm) at BBS, as detected by α-RRxS/T-p antibody, is higher than WT, with Rim4(5C) and Rim4(PxLm+5C) as negative controls. C) The presentative FM images showing the intracellular distribution of Rim4 variants with EGFP tagged at N-terminus (green), including Rim4, Rim4(PxLm), and Rim4(PxLm+5C); Nup49-mScarlet marks the nuclear membrane (red). The arrow (white) indicates the nucleus. D) The sporulation efficiency of indicated Rim4 variants, presented as the tetrad percentage, is associated with (C). More than 300 cells were counted for each strain. E) Top, Schematic of Rim4(PxLm) and Rim4(R6-A) mutants. Bottom, the presentative FM images show the subcellular distribution of EGFP-tagged Rim4(PxLm) and Rim4(R6-A) mutants. The arrow (white) indicates the nucleus. R6-A reduced the nuclear retention of Rim4(PxLm). F) Top, Schematic of Rim4 variants created to rescue Rim4(PxLm), including Rim4(SA_1_), Rim4(SA_2_) and Rim4(SA_1/2_). Bottom, the presentative FM images show the subcellular distribution of indicated Rim4 variants. The arrow (white) indicates the nucleus. SA_1_ and SA_1+2_ partially reduced the nuclear retention of Rim4(PxLm).

Next, we examined whether the S/T-p residues at the BBSs are primary targets of Cdc14 on Rim4. We predicted that Rim4(5C) should rescue the nuclear retention phenotype in Rim4(PxLm) if they are. Remarkably, FM analysis of the Rim4(PxLm+5C) mutant strain confirmed our prediction (Fig. 5C). In contrast, Rim4(5C) failed to rescue the subcellular distribution of Rim4(FLm), which contains terminally dead RRM-1 and RRM-3 due to disrupted RNA binding pocket (Fig. S5A). Our data demonstrate that Rim4(PxLm) has intact mRNA binding ability but lacks a trigger, i.e., Cdc14-mediated phosphorylation, to form a complex with mRNAs. Although Rim4(PxLm+5C) restored WT-like subcellular distribution, to our surprise, the 5C mutation failed to rescue sporulation in Rim4(PxLm) cells (Fig. 5D). This result highlights the importance of temporal Rim4-Bmh1/2 interaction, meanwhile, inspiring us to examine BBS-independent P-sites on Rim4, as additional Cdc14 targets.

### Cdc14 physiologically de-phosphorylates Rim4 beyond the BBSs

The Cdc14 docking site, i.e., P_454_E_455_L_456_, resides in the R6, conveniently positioned Cdc14 to the 11 S/T residues in this region. Earlier, we showed that R6-E exhibited nuclear retention (Fig. 1E). Therefore, we hypothesize that Cdc14 also mediates de-phosphorylation in the R6 region. Indeed, combining R6-A with PxLm mutants enabled a relatively balanced subcellular distribution of Rim4 (Fig. 5E), albeit not as effective as the 5C mutant (Fig. 5C), indicating that R6 harbors P-sites targeted by Cdc14, facilitating Rim4 nuclear export.

Cdc14 prefers a Serine-Proline (SP) target, especially in an unstructured region^31^–^33^. Rim4 contains two such SP motifs in its LCD, i.e., S_532_P_533_ (SP_1_) and S_658_P_659_ (SP_2_) (Fig. 5F, top). By mutating S_658_ into Alanine (S_658_A, i.e., SA_2_), the nuclear retention of Rim4(PxLm) was moderately reduced, while mutating S_532_ into alanine (S_532_A, i.e., SA_1_) has little effect (Fig. 5F). Like R6-A mutant, the effect of SA_2_ on Rim4(PxLm) distribution is less penetrating compared to the 5C mutation (Fig. 5C). Nonetheless, SA_2_ and R6-A slightly increase the sporulation efficiency of Rim4(PxLm) (Fig. S5B), indicating that S_658_P_659_- (SP_2_) and some S/T residues yet-to-be-determined in R6 region are physiological P-sites handled by Cdc14. In summary, Cdc14 coordinately de-phosphorylates Rim4 at multiple P-sites to stimulate Rim4 nuclear export, presumably by assisting Rim4-mRNA assembly.

### Cdc14 sensitizes the Rim4 degradation by autophagy

Thus far, our data indicate that Cdc14 dissembles the Rim4-Bmh1/2 complex in the nucleus, leading to Rim4-mRNA assembly. However, during the meiotic divisions, Cdc14 at anaphases temporally resides in the cytoplasm^34^. So, what would be the consequence if Cdc14 triggers Rim4-Bmh1/2 disassembly in the cytoplasm, where autophagy is active while the mRNA level is low?

Previously, we found that Rim4 forms cytosolic puncta that co-localizes with Atg8, an indication of the engulfment of Rim4 by autophagosomes during meiotic divisions (accompanying manuscript). Remarkably, FM analysis of Rim4 (mScarlet-Rim4) puncta detected no co-localization with Bmh1/2 (Bmh1/2-EGFP) (Fig. 6A). Therefore, Bmh1/2 is not in complex with the Rim4 degraded by autophagy. Next, we adapted a chemical genetic approach (Fig. S6A) to measure Atg1-as (Atg1[M102G]) activity *in vitro*^33^, which can be stimulated by some substrates of selective autophagy to initiate their autophagic degradation^35,36^. As a substrate of selective autophagy, recombinant Rim4 and Rim4_BBS_-p (by PKA treatment) indeed stimulated Atg1-as at a concentration of ^~^0.2 to ^~^1.0 μM, a physiological range of intracellular Rim4 (Fig. 6B; S6B). Remarkably, Bmh1/2 suppressed Rim4_BBS_-p-triggered Atg1 activation (Fig. 6B). We noticed that Rim4_BBS_-p alone, at low concentration (0.2 μM), is less efficient than Rim4 in activating Atg1 in the cell lysate. We interpret that it is due to the free endogenous Bmh1/2 in the cell lysate forming complexes with a portion of supplemented Rim4_BBS_-p. In line with this observation, next, we showed that immunoprecipitated Rim4-FLAG complex (IP: α-FLAG) from prophase I cell lysates (Fig. 2A; lane 1), a stage before Cdc14 relocation to the cytoplasm and Rim4 clearance, suppressed Atg1-as activity (Fig. S6C). Thus, we concluded that Bmh1/2 protects Rim4 from autophagy in the cytoplasm.

**Figure 6.**
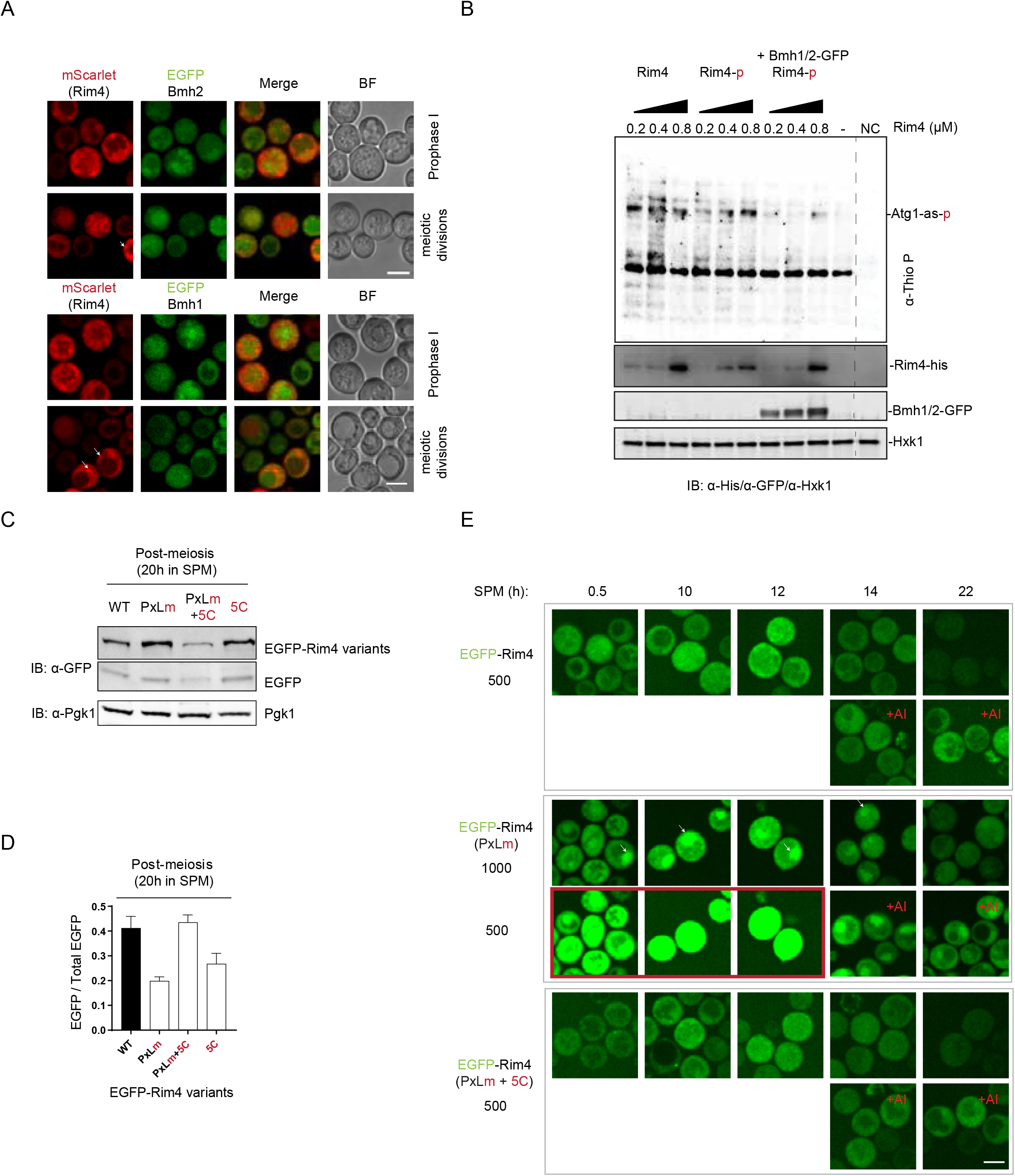
Cdc14 sensitizes the Rim4 degradation by autophagy. A) The presentative FM images showing the subcellular distribution of mScarlet-Rim4 (red) and EGFP tagged Bmh2 (top panel) or Bmh1 (bottom panel) (green). The arrow (white) indicates the Rim4 puncta that do not co-localize with Bmh1/2. Images are selected from 3 independent experiments with over 1,000 cells from each condition analyzed. Prophase I cells are collected at 12 hr in SPM. Meiotic division cells were collected at 14h and 16h. The scale bar is 5 μm. B) IB analysis of Atg1-as kinase activity in prophase I cell lysates, which were supplemented with Rim4, PKA-phosphorylated Rim4 (Rim4-p) or the mixture of Rim4-p and BMH1/2-EGFP, as indicated. The Atg1 kinase assay was as described in Fig. S6A. Hxk1, hexokinase isoenzyme 1 (loading control); Atg1-as-p, Atg1-as auto-phosphorylation. -, no Rim4 supplemented; NC, no supplemented ATP in cell lysate. C and D) Shown are (C) images and (D) quantification of IB analysis of indicated Rim4 variants in post-meiosis (20h in SPM) cell lysates using an α-GFP antibody. In (D), the EGFP process, an indication of autophagic degradation of EGFP-Rim4, was measured by dividing the free EGFP by the total EGFP signal. E) The presentative FM images of indicated EGFP-Rim4 variants at different time points in SPM, with (+AI) or without 5μM 1NM-PP1 applied at prophase I (12 hr in SPM). 1NM-PP1 specially inhibits Atg1-as (Atg1-M_102_G) and hence autophagy. To show the subcellular texture of Rim4 variants, the maximum display value for each strain was adjusted as depicted. The scale bar is 5 μm.

Intriguingly, the purified Rim4-FLAG protein complex stimulated Atg1-as activity after Cdc14 treatment (Fig. S6C), presumably due to the loss of Rim4-Bmh1/2 interaction triggered by Cdc14. Consistent with this scenario, the Cdc14-binding defective Rim4(PxLm) is more stable than Rim4 and resistant to autophagy during meiotic divisions (Fig. 6C; 6D). By mutating the BBSs to disrupt Bmh1/2 binding, the resulting Rim4(PxLm+5C) was less stable than Rim4 and restored its autophagic degradation (Fig. 6C; 6D), indicating that Cdc14-mediated Bmh1/2 dissociation helps autophagic Rim4 degradation during meiotic divisions. Notably, mRNAs stabilize Rim4 in the RNPs (accompanying manuscript). Rim4(5C) exhibited enhanced nuclear export, presumably due to its inability to bind Bmh1/2, thereby binding better with mRNAs (Fig. 3A). Therefore, it is not surprising that Rim4(5C) is more stable than Rim4 (Fig. 6C; 6D).

Lastly, using FM, we showed that Rim4 (EGFP-Rim4) cellular level decreased during *NDT80*-synchronized meiotic divisions in an autophagy-dependent manner (Fig 6E), as we previously reported^17^. Consistent with IB analysis (Fig. 6C), the cellular level of Rim4 (PxLm) is generally higher than WT Rim4 from meiosis entry until the end of meiosis (Fig. 6E). Remarkably, the nuclear retention phenotype of Rim4(PxLm) gradually disappeared upon *NDT80* expression (Fig. 6E). This phenomenon further supports that Cdc14 facilitates Rim4-mRNA assembly in the nucleus; due to its relocation into the cytoplasm at anaphases, Cdc14 lost its influence on Rim4 (Rim4-mRNA) subcellular distribution. Strikingly, inhibition of autophagy during meiotic divisions protects the nuclear retention of Rim4(PxLm) (Fig. 6E), indicating that autophagy-mediated Rim4 degradation prevents Rim4 from entering the nucleus to bind mRNAs again. Therefore, autophagy is required to effectively release and translate Rim4-sequestered mRNAs, supporting our previous finding that *CLB3* translation needs active autophagy^17^.

In summary, our findings support a model that spatiotemporally controlled phosphorylation and de-phosphorylation determine Rim4 function and protein stability (Fig. S6E). Before meiotic divisions, in the nucleus, the cell cycle phosphatase Cdc14 dephosphorylates Rim4, resulting in Bmh1/2 dissociation from Rim4. Because Bmh1/2 acts as an mRNA competitor on Rim4, the release of Bmh1/2 facilitates Rim4-mRNA RNP formation, suppressing translation. During meiotic divisions, we propose phosphorylation (by PKA) drives the transition from Rim4-mRNA to Rim4-Bmh1/2, releasing mRNAs for translation. Next, Cdc14 relocates to the cytoplasm and triggers Rim4-Bmh1/2 disassembly and autophagic Rim4 degradation while stimulating autophagy^33^. Thus, PKA and Cdc14 coordinately control the timing of autophagic Rim4 degradation and couple it with the translation of Rim4 target mRNAs (Fig. S6E).

## Discussion

### The roles of Bmh1/2 interaction with Rim4

The effect of 14-3-3 (Bmh1/2 in yeast) docking on a phosphorylated protein can vary. With its rigid α-helical backbone, the 14-3-3 proteins can alter the structure of their binding partners^37^. Strikingly, we found no other macromolecular components, such as RNAs or proteins, in the soluble heterotrimeric Rim4-Bmh1/2 complex isolated from meiotic prophase I cell lysates. Therefore, we think Bmh1/2 proteins, which are abundantly presented in both nucleus and the cytoplasm compartment and affect a broad range of biological processes^23,38–43^, serves as a sequester of phosphorylated Rim4 (Rim4_BBS_-p).

Rim4 harbors four putative Bmh1/2 binding sites; at least three have been experimentally verified in this study, including one in the RRM2 (T_216_). Our data demonstrate that Rim4 and mRNA compete on Rim4 binding, presumably in the Bmh1/2 binding sites (BBS)-containing RRMs, or unstructured segments that regulate Rim4-mRNA interaction as computationally predicted (Fig. S2B; S2C)^44^. Depending on the binding affinities yet-to-be-determined, Bmh1/2 either keeps mRNAs away from Rim4 or generates a threshold for Rim4-mRNA complex formation^43,45,46^. Either way, Bmh1/2 binding delays Rim4 nuclear export by inhibiting Rim4-mRNA complex assembly. We speculate that this mechanism ensures Rim4 interacts only with its *bona fide* mRNA substrates that tightly associate with the three RRMs of Rim4.

Both Bmh1/2 and mRNAs protect Rim4 from autophagy. The fact that mRNAs and Bmh1/2 share common binding sites on Rim4 implies that mRNAs and Bmh1/2 protect Rim4 by one mechanism. Rim4 was selectively degraded by autophagy in an Atg11-dependent manner in the absence of mRNAs (accompanying manuscript). Bmh1/2 and mRNAs may mask the same sites on Rim4 that engage autophagy machinery, e.g., Atg11 or Atg8^47,48^. When the time comes for the translation, PKA phosphorylates Rim4; next, Bmh1/2 facilitates timely mRNA release from phosphorylated Rim4. Then, Cdc14-triggered Bmh1/2 dissociation stimulates Rim4 degradation by autophagy, thereby preventing Rim4 from entering the nucleus to rebind mRNAs. Thus, Rim4 is only efficiently degraded by autophagy free of mRNAs and Bmh1/2, requiring sequential PKA-mediated phosphorylation and Cdc14-mediated de-phosphorylation in the cytoplasm.

14-3-3 proteins can regulate the nuclear import of their substrates^49^–^52^. Rim4 harbors multiple BBSs. Interestingly, the occupation of one BBS, i.e., the site of S_607_, by Bmh1/2 will mask a nearby NLS of Rim4 that contains K_600_R_601_, Identified in our accompanying manuscript.

Likely, Bmh1/2 will delay the nuclear import of Rim4 by masking the NLS via phosphorylated S_607_, *e.g*., combined with phosphorylated S_525_ (Fig. S2B; state III). Thus, Bmh1/2 binding might affect the nucleus-cytoplasm transport of Rim4 bi-directionally, depending on which BBSs are phosphorylated, thereby assisting a balanced subcellular Rim4 distribution. This observation implies why S_525_C and S607C can partially rescue the reduced sporulation of T_216_C, S_367_C and T_368_C, i.e., in the Rim4(5C) strain (Fig. 2G). We speculate that Bmh1/2 can affect various meiotic Rim4 activities in responding to the meiotic dynamics, e.g., Rim4-mRNA assembly and disassembly, Rim4-autophagy interaction, and Rim4 nuclear import, by binding to different pairs of BBSs.

### PKA-mediated site-specific phosphorylation on Rim4 versus Ime2-mediated cumulative phosphorylation

The kinase and phosphatase coordinately control 14-3-3 interaction with their partners^25,53^. To date, Ime2 is the only identified kinase for Rim4^12,54^. We found that PKA can specifically phosphorylate the BBSs of Rim4, while Ime2 does not (Fig. 3B; S3A); consistently, the sequences of Rim4 BBSs share a classical consensus of PKA target, i.e., RRxS/T-p. Moreover, Ime2-mediated cumulative phosphorylation of Rim4 stimulates Rim4 clearance via proteasome-mediated degradation during meiotic divisions^15^, while PKA-mediated Rim4 phosphorylation at the BBSs peaks before meiotic divisions (Fig. 3D) and stabilizes Rim4 against autophagy.

Interestingly, S. G. Herod et al. previously reported that Bmh1/2 could stimulate Ime2-mediated Rim4 phosphorylation, thereby facilitating proteasome-mediated Rim4 degradation^16^. Therefore, it is likely that PKA and Ime2 modulate Rim4 at distinct P-sites and at different meiotic stages, engaging different effectors. Furthermore, Cdc14, which de-phosphorylates Rim4, generally antagonizes the cyclin-dependent kinase CDK1 activities^53,55,56^; therefore, CDK1 (Cdc28 in yeast) may also be involved in Rim4 phosphorylation. Knowing the total kinase and phosphatase network of Rim4 modification is vital for further dissection of PTM-modulated Rim4 meiotic functions.

The most critical Rim4 functional interactor is mRNA^13,17,21^. Rim4 harbors three structured RRM domains to its N-terminus to accommodate mRNAs. Notably, the T_216_-, S_367_/ T_368_-containing BBSs are part of the RRMs. As a result, phosphorylation states of T_216_ and S_367_/ T_368_ will logically regulate Bmh1/2 interaction with RRM2, thereby affecting Rim4-mRNA complex formation. It is unknown, though, whether phosphorylation at the RRMs can influence Rim4-mRNA interaction independent of PKA and Bmh1/2. Interestingly, phosphorylation and de-phosphorylation mimicking mutants have distinct effects on Rim4’s three RNA-binding RRMs. For instance, Rim4-R2A (RRM1) and Rim4-R3A (RRM2), but not Rim4-R2E nor Rim4-R3E, showed nuclear retention. This finding indicates that phosphorylation at RRMs and flanking sequences could modulate the Rim4-mRNA complex assembly^57,58^.

In addition, beyond the RRM domains, previously, LCD-mediated Rim4 self-assembly was found to enhance Rim4-mRNA interaction^15,59,60^. Therefore, de-phosphorylation may stimulate Rim4 self-assembly upon Bmh1/2 dissociation, enhancing Rim4-mRNA interaction. Vice versa, phosphorylation-mediated Rim4 disassembly on its own might reduce Rim4-mRNA affinity.

### The dual role of Cdc14 in Rim4 regulation

Cdc14 has a structurally defined binding preference for a PxL motif^30^, identified in Rim4 and characterized in this study. Cdc14 resides in the nucleus except for the anaphases^34,61^. At early meiotic stages, we found that Rim4 dynamically shuttles between the nucleus and the cytoplasm compartments, requiring Cdc14 to facilitate its nuclear export. Our data demonstrate that Cdc14 de-phosphorylate Rim4 dissociates its bound Bmh1/2 in the nucleus and the cytoplasm. However, the nucleus and the cytoplasm compartments differ regarding mRNA and autophagy availability; the former actively produces the mRNAs but lacks autophagy machinery. Therefore, the consequences of Rim4-Bmh1/2 disassembly in the two subcellular compartments also differ: Bmh1/2-free Rim4 forms a complex with mRNAs in the nucleus and remains stable, whereas being degraded by autophagy in the cytoplasm, determined by the subcellular localization of Cdc14.

Notably, we previously reported that Cdc14 docks to PAS, the autophagy initiation sites, and stimulates autophagy initiation by de-phosphorylating Atg13 to activate Atg1 upon being relocated into the cytoplasm during meiotic anaphases^33^. Therefore, Cdc14, after its relocation to the cytoplasm, spatiotemporally couples upregulated autophagy to Rim4 degradation. Lastly, other than Rim4, we speculate that Cdc14 might have a general role in RNP assembly in the nucleus, and regulate the degradation of a list of RBPs by autophagy during meiotic divisions, thereby influencing meiotic translation post-transcriptionally. To this end, investigating whether these RBPs in complex with 14-3-3 proteins controlled by Cdc14 activity would be an important first step.

**Figure S1. Phosphorylation regulates the subcellular distribution of Rim4**

A) Schematic of induced expression of *NDT80* (*pGAL1-NDT80*), for synchronized metaphase I entry, by 1 μM β-estradiol. 1 μM β-estradiol was added to meiotic cells after 12 h in SPM to synchronize the entry of meiotic divisions.

B) FM images showing the subcellular distribution of EGFP-Rim4 variants, i.e., WT and the indicated mutations (R1-A to R9-A; R1-E to R9-E) (green), in cells arrested at prophase I (GAL-NDT80 system). Nuclear membranes are marked by Nup49-mScarlet (red). Due to their higher subcellular levels, the maximum display value of EGFP-Rim4(R9-A) and EGFP-Rim4 (R8-E) is set to 1000 and 1500, respectively, while the others are set to 500.

C) Prediction of Rim4 disordered regions with the DISOPRED3 program^22^. Precision is the ratio of true positive residues number divided by total positive residues number, a score reflecting the disorder level of the local area; the higher the score is, the more disorder the position is.

**Figure S2. The schematics of Bmh1 and Bmh2 chaperoning phosphorylated Rim4**

A) The Rim4 protein structure was predicted by AlphaFold^62^, and the RRM-2 and RRM-3 domains of Rim4 were separately predicted by Robetta^63–65^. The RRMs are highlighted in orange; predicted BBSs are marked in dark green; The shadows show the predicted electrostatic. The positively charged areas are marked with red, the negatively charged areas are marked with blue, and the neutral areas are marked with white.

B) Model of possible Bmh1/2-Rim4 complexes as a response to different phosphorylation states of Rim4 at the BBSs. State II is modeled as an intermediate state of state I and state II, with S_525_ site unchanged.

C) Schematic of the overlap of Bmh1/2 binding sites (S_525_, S_607_) with RNA interaction motifs predicted by ProNA2020^44^. The blue box indicates RNA binding sites with a reliability index (RI) between 0 and 33; the red box with RI between 34 and 66.

**Figure S3. PKA, but not Ime2, phosphorylates Rim4 in vitro at the BBSs**

A) PKA catalytic subunit (Sigma, P2645) mediated phosphorylation of recombinant Rim4-6×His at the RRxS/T sites was conducted *in vitro*, compared with Ime2 treatment. Shown are IB analyses of the proteins after 1h of kinase treatment at room temperature, using an α-RRx-S/T-p antibody that specifically recognizes phosphorylated RRx-S/T and α-His to show total Rim4.

B) The Ponceau S staining of the blot in Fig. 3F. Bmh1-FLAG (*pBMH1-Bmh1-FLAG*) co-immunoprecipitation with Bmh1 and Bmh2, as well as with its phosphorylated substrates (Fig. 3F).

C) Coomassie blue staining image showing the purified recombinant proteins from *E. Coli*.

**Figure S4. The onset of the nuclear retention of Rim4(PxLm) is before prophase I**

A) FM images showing the subcellular distribution of EGFP-Rim4(PxLm) over time in SPM.

**Figure S5. The effects of Rim4 mutagenesis and *in vitro* treatment**

A) Presentative FM images of the subcellular distribution of EGFP-Rim4(FLm) or EGFP-Rim4(FLm+5C) (green). Nuclei are marked by Nup49-mScarlet (red).

B) The sporulation efficiency (presented by tetrads percentage) of strains expressing EGFP-Rim4 variants, including WT, PxLm, R6-A, PxLm+R6-A, SA_1/2_, PxLm+SA_1_, PxLm+SA_2_, PxLm+SA_1/2_, and PxLm+5C. More than 300 cells were counted for each strain.

**Figure S6. Cdc14 de-phosphorylated Rim4 meets different fates in the nucleus and the cytosol**

A) Schematic of the chemical-genetic strategy for monitoring Atg1-as (analog-sensitive Atg1, Atg1-M_102_G) kinase activity in vitro. Atg1-as thiophosphorylates its substrates with a bulky ATPγS analog (N6-PhEt-ATP-γ-S). Thiophosphorylated substrates of Atg1-as can then be alkylated with para-nitrobenzyl mesylate (PNBM) and detected by IB using anti-thiophosphate ester (α-thioP) antibodies.

B) The measurement of intracellular Rim4 concentration with recombinant Rim4-EGFP as standard by IB. The signals of Rim4-EGFP in the cell lysates and the recombinant Rim4-EGFP standards were plotted on an X-Y coordinate, fitted with a simple linear regression equation, to calculate the intracellular concentration. See the method part for more details.

C) IB analysis of Atg1-as kinase activity in prophase I cell lysates. Immunoprecipitated FLAG-Rim4-BMH1/2 complex (Fig. 2A, 1^st^ lane), treated by Cdc14 or mock treatment, was supplemented to the cell lysates before Atg1-as kinase assay. The whole-cell lysates were then subjected to IB with indicated antibodies. Hxk1, hexokinase isoenzyme 1 (loading control); Atg1-as-p, Atg1-as auto-phosphorylation. Minus (-), mock elute (Fig. 2A, 2^nd^ lane) was supplemented to the cell lysates before Atg1-as kinase assay. NC, no supplemented ATP in cell lysate.

D) The model of Cdc14 spatiotemporally regulating Rim4 function and stability. Notably, the cell cycle phosphatase Cdc14-mediated Rim4-Bmh1/2 disassembly stimulates Rim4-mRNA complex formation and autophagic Rim4 degradation in the nucleus and the cytoplasm, respectively. Before meiotic divisions, Cdc14 mainly stimulates Rim4-mRNA assembly in the nucleus (left); the subcellular relocation of Cdc14 from the nucleus to the cytoplasm during meiotic anaphases, coordinated by PKA controls the timing of Rim4 degradation, thereby allowing translation of Rim4-sequestered mRNAs (right).

## Supporting information

Key Resources Table

