## Supplementary figures and images for "Cdc14 spatiotemporally regulates Rim4-mRNA complex assembly and stability during meiosis"

### Figure S1

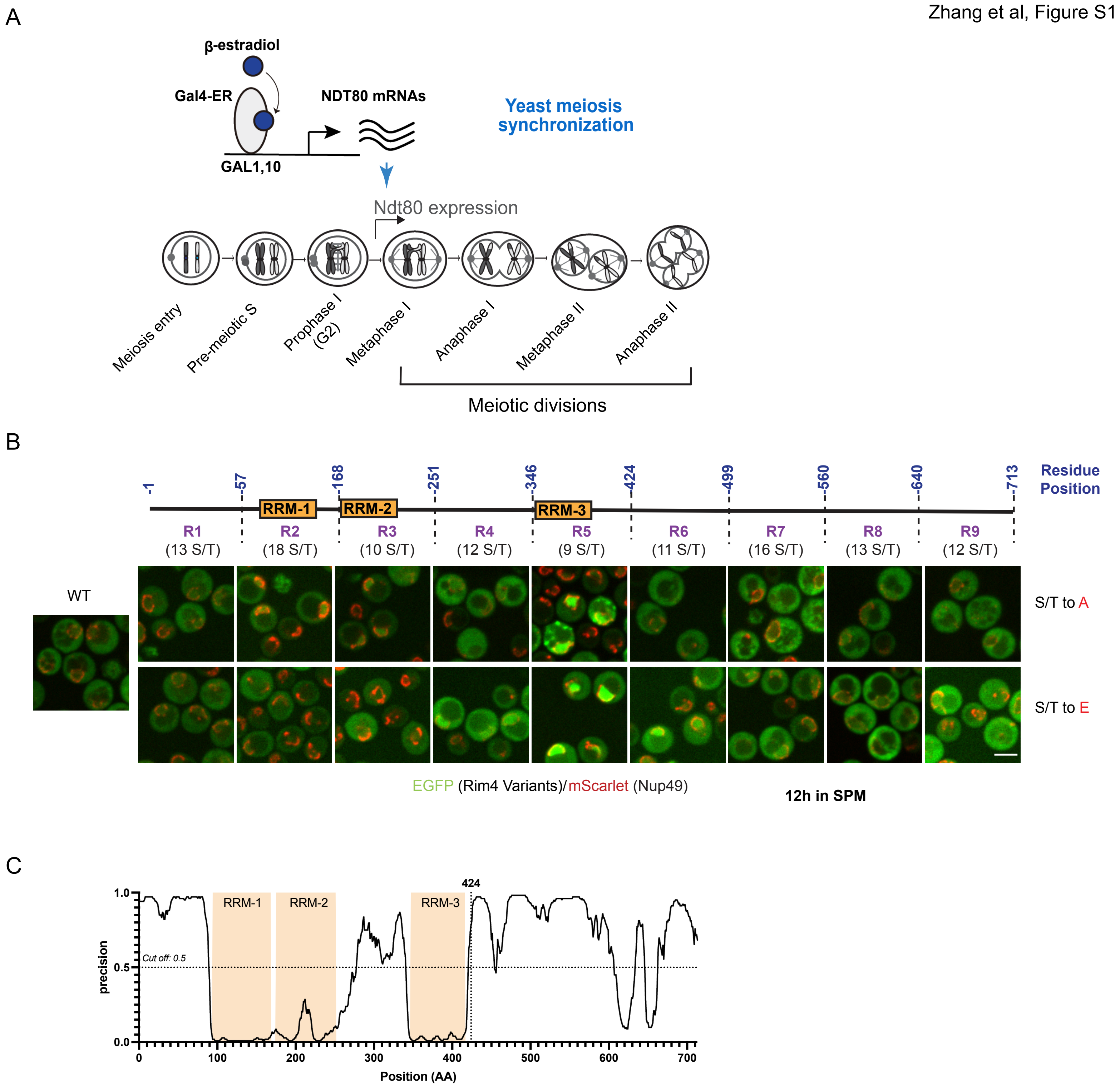

### Figure S2

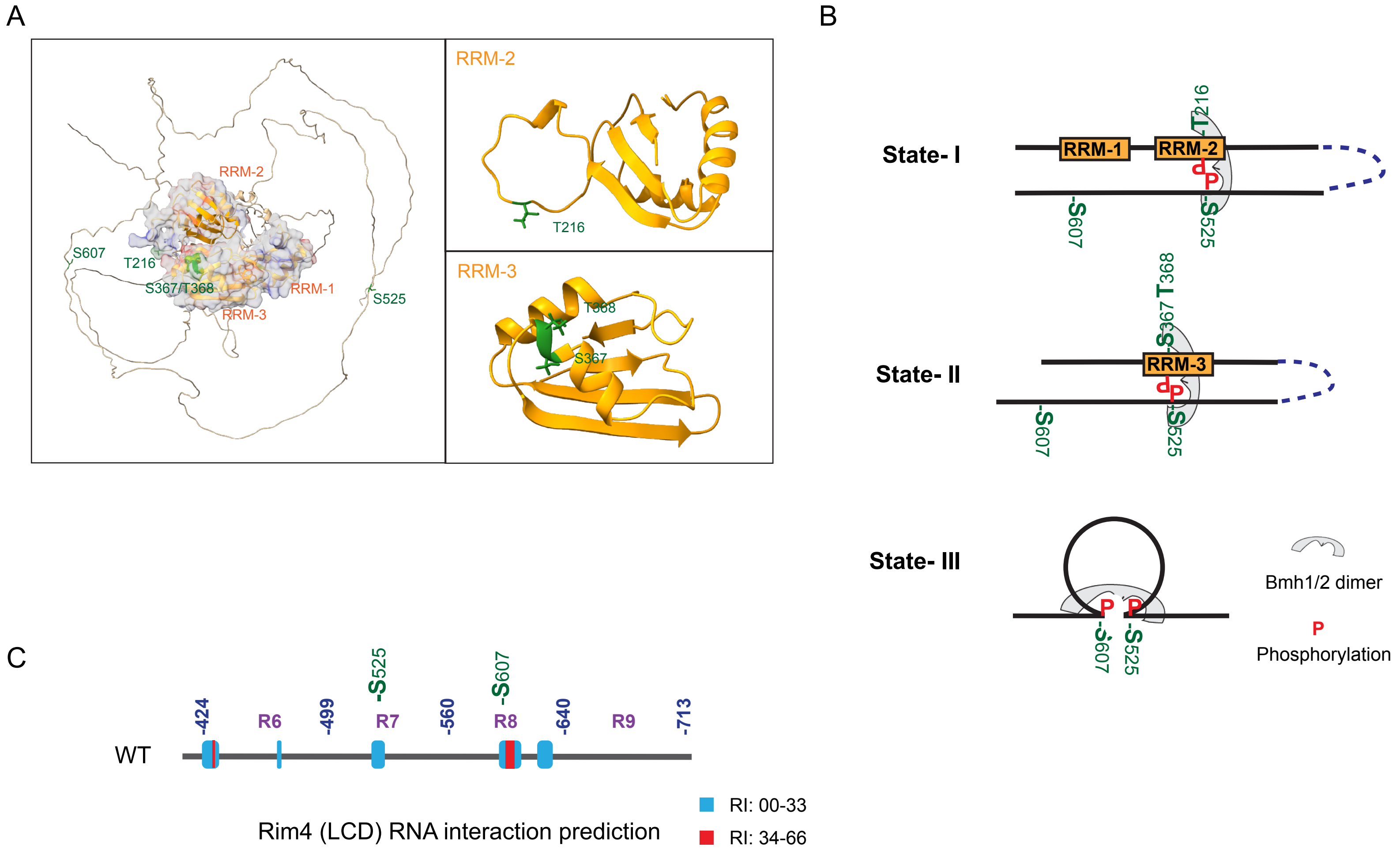

### Figure S3

A

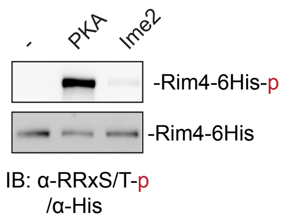


B

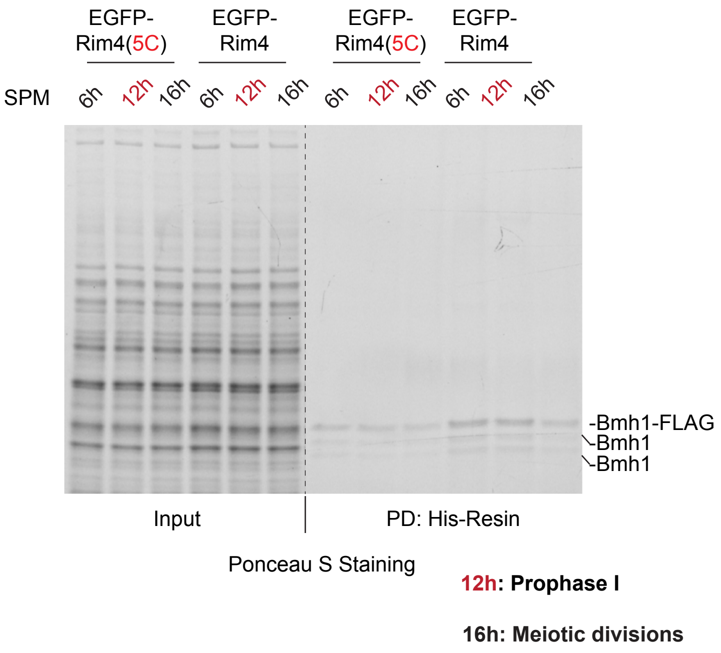


C

Recombinant proteins in this study

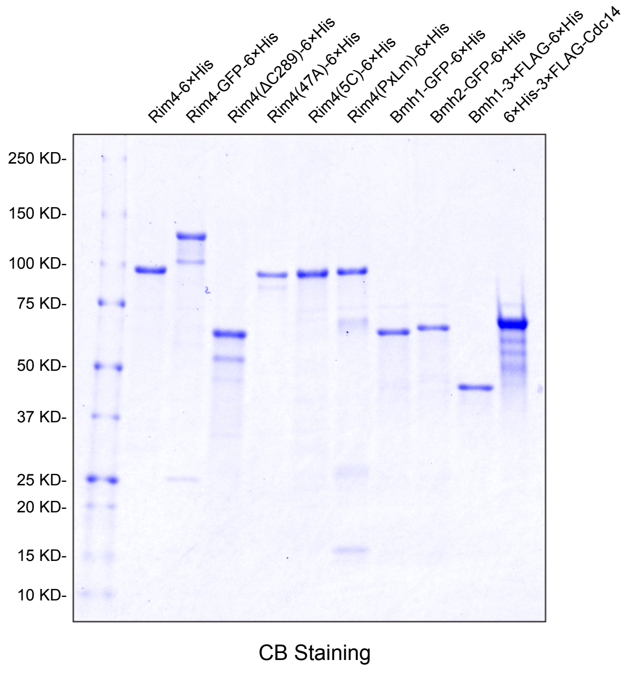

### Figure S4

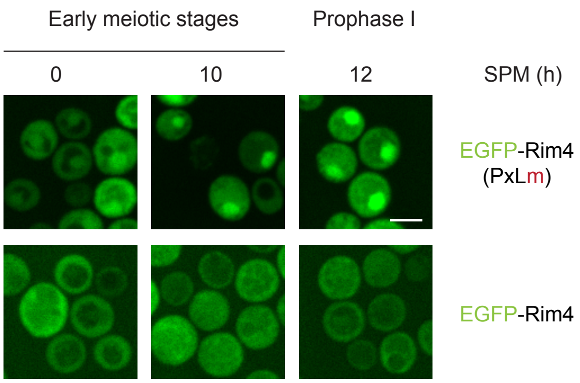

### Figure S5

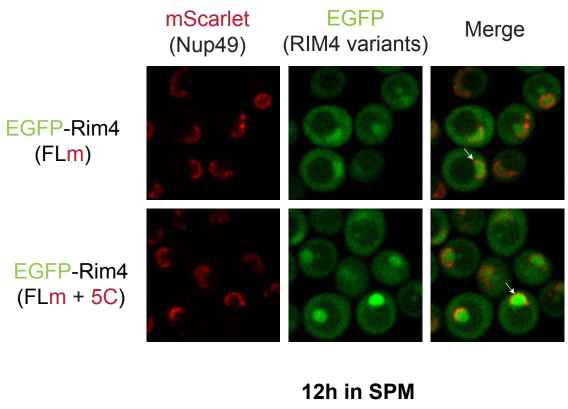

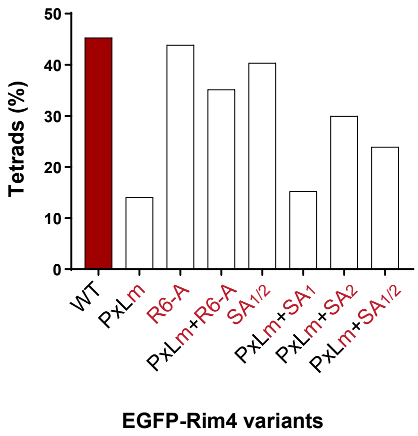

### Figure S6

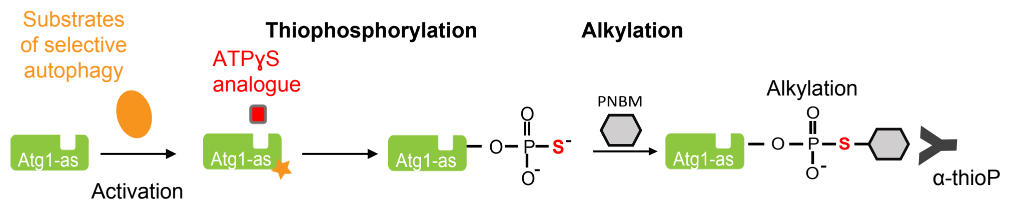

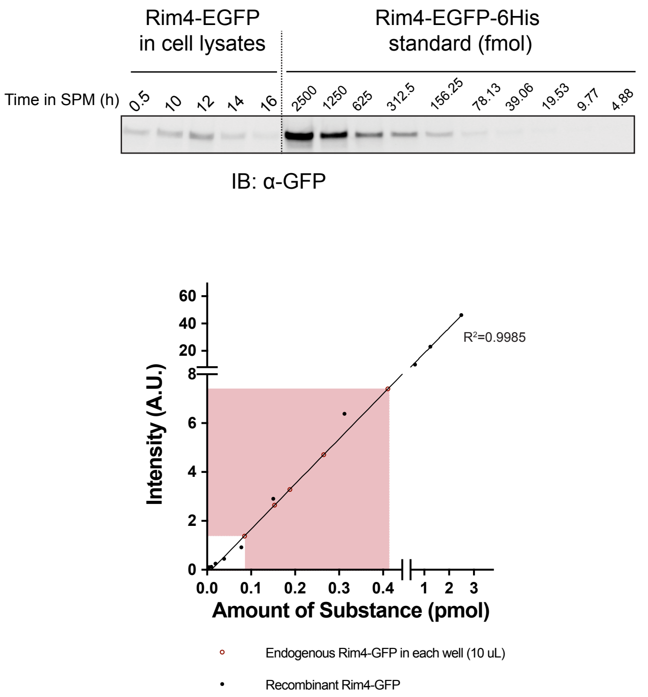

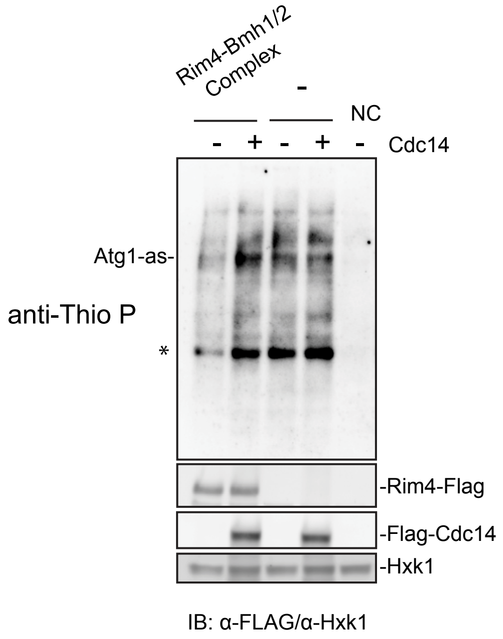

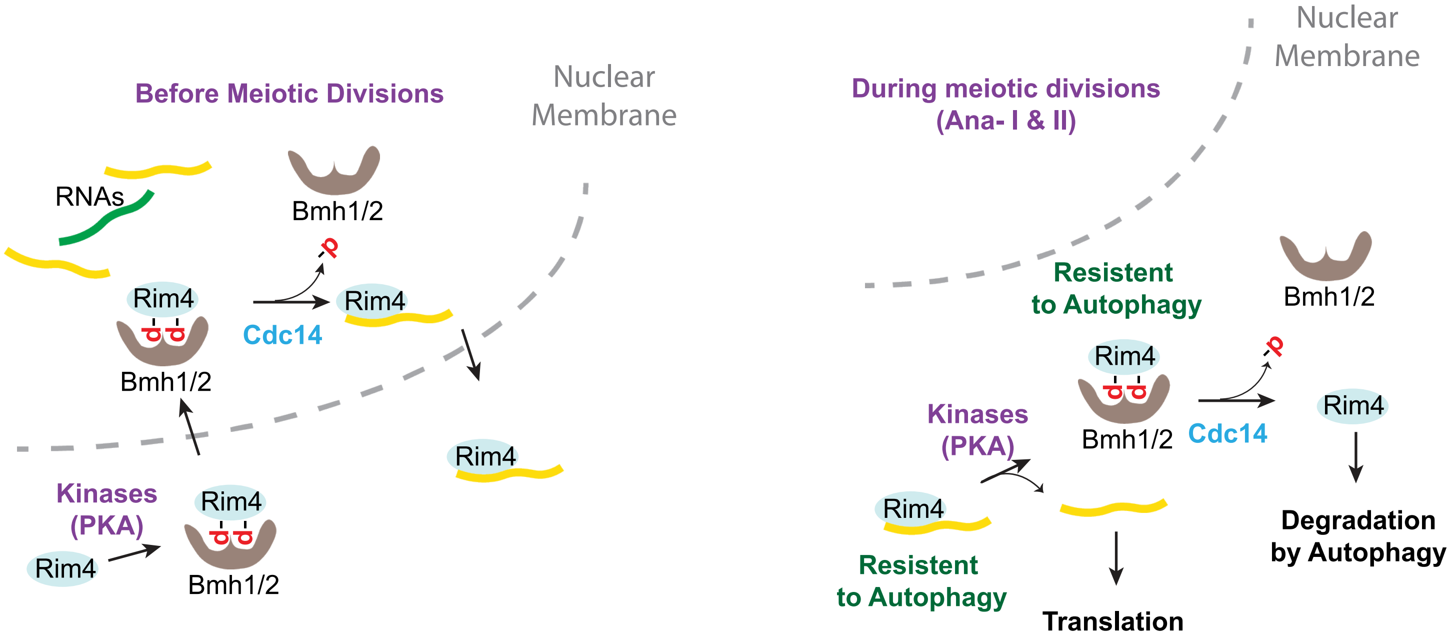
