## Supplementary material for "Cdc14 spatiotemporally regulates Rim4-mRNA complex assembly and stability during meiosis": Methods

**STAR Methods**

**Experimental Model and Subject Detains**

**S. cerevisiae Strain and Manipulation**

Saccharomyces cerevisiae strains were derivatives of W303 (ade2-1 his3-11,15 leu2-3,112 trp1-1 ura3-1 can1-100) (Table S1). Unless otherwise indicated, the analog sensitive version of ATG1(Atg1-as) was created by gatekeeper residue change (M_102_G) as described earlier^1,2^ and introduced into the background of listed strains to allow conditional autophagy inhibition by 1NM-PP1.

Deletion strains were constructed in a parent background by PCR-mediated knock-out with one of the following drug resistance or prototrophic marker cassettes: pFA6a-*kanMX6*/pFA6a-*NatMX*^3^ /pKlURA/pCgHIS^4^. C- terminal tagging at the endogenous genomic locus was introduced by PCR-mediated epitope tagging as previously described^5^.

Specifically, strains with GAL-NDT80 GAL4-ER were constructed by replacing the endogenous NDT80 promoter with the inducible GAL1,10 promoter as described^6^.

The N-terminus EGFP tagged Rim4 strains were created by the following steps: the endogenous RIM4 ORF was deleted by URA3 as described above. Then the pRS303 backbone plasmids carrying *pRIM4:EGFP-Rim4* *variants* (See Table S2) were linearized and integrated into the ∆rim4 yeast genome at his3 locus and selected by histone prototrophy, respectively.

The *pNUP49:Nup49-mScarlet* fragment was carried on a pRS304 backbone plasmid (pFW338). The plasmid was linearized and integrated to the yeast genome at trp1 locus, and selected by tryptophan prototrophy.

All strains used in this study were diploids mated by the haploids harboring the needed genotype (Table S1).

**Media**

The following media were used in this study. YPD (2% peptone, 1% yeast extract, 2% glucose), YPA (2% peptone, 1% yeast extract, 2% potassium acetate), SD (0.67% yeast nitrogen base, 2% glucose, auxotrophic amino acids and vitamins), and standard sporulation medium SPM (0.6% potassium acetate, pH 8.5). The SD-dropout medium was made with dropout stock powder lacking histidine (-His), tryptophan (-Trp), leucine (-Leu) and/or uracil (-Ura).

**Method Details**

**Sporulation, Vegetative Growth, and Synchronized Ndt80 Arrest/Release**

The cells were inoculated from a single colony of a fresh YPD streak to YPD medium, and cultured at 30˚C with 250 rpm shaking for overnight. Backdiluted the strain to fresh YPD medium to concentration of 0.2 OD_600_. The cells were then growing at 30˚C with 250 rpm shaking. The cells reached to about 0.8 OD_600_ after 4 hrs culture, and the cells were collected for transforming.

A single colony of yeast strains from YPD plate were picked, spread on YPG (3% glycerol) plates and grown at 30°C for 2 days. Colonies grown out were spread to YPD plates and grown until cells formed a lawn (~ 24 hours). Cells on plates were collected and suspended in YPA liquid medium (OD_600_=0.3) and grown for 14-16h at 30°C. Cells were then pelleted, washed with water twice and resuspended in SPM to final OD_600_=2. For M-phase synchronization, following incubation in SPM for 12 hours, the synchronized strains containing *pGAL1:Ndt80* and *GAL4(848)-ER* recombinant transcription regulator were released from the prophase I arrest by addition of 1 μM β-estradiol to induce Ndt80 expression. To assess the sporulation efficiency, percentage of tetrads were counted after 48 hours in SPM at room temperature. At least 300 cells for each strain were counted under bright field with Olympus microscope (BX40, 40x objective).

**DNA Replication Analysis with Flowcytometry**

Cells were collected by 12 hr and fixed in 70% ethanol at 4˚C for overnight. Afterwards, the cells were pelleted and resuspended in 50 mM Sodium Citrate (pH 7.0) followed by sonication at 30% power for 15 sec. Cells were pelleted again and resuspended in the same solution. The cells were treated with 0.25 mg/mL RNase A at 37˚C for overnight. Next, the cells were incubated in 1 µM SYTOX Green dye at RT in the dark for at least 1 hr before analyzed on flow cytometer (BD FACSCalibur™). The forward scatter (FSC) and side scatter (SSC) were used to gate the living cells. The cell counts of green (530/30 nm) signal in gate were collected and analyzed by Flowing Software 2.5.1 (<https://bioscience.fi/services/cell-imaging/flowing-software/>).

**Plasmid Construction**

Plasmids used in this study are listed in Table S2. pRS303-*pRIM4:EGFP-Rim4* (pFW208) was constructed by subcloning genomic Rim4 with its promoter and terminator amplified by PCR into SalI/NotI restriction sites of pRS303, followed by inserting EGFP with a linker containing a PacI restriction site amplified by PCR between the promoter and start codon of Rim4 by Gibson Assembly^6^. The mutagenesis of Rim4 as shown in Table S2 were introduced by overlap extension PCR to amplify the mutant Rim4 cds and terminator and cloned into PacI and NotI of pFW208.

The plasmids for Rim4 variants expression were constructed by subcloning PCR-amplified Rim4 or Rim4 mutants CDS into the NdeI and KpnI restriction site of pET29b(+) by Gibson Assembly. The plasmids for Bmh1/2-GFP and Bmh1-3FLAG expression were constructed by subcloning overlap extension PCR-amplified Bmh1/2-GFP or Bmh1-3FLAG into NdeI and KpnI restriction sites of pET29b(+).

**Protein Expression and Purification**

**Recombinant protein purification**

pET-based protein expression in BL21 (DE3) E. coli cells was induced by IPTG as described previously (Wang et al., 2010). RIM4 variants-6×His/6×His-3Flag-Cdc14 were expressed at 16°C overnight. Cells were collected by centrifugation, resuspended in bacteria lysate buffer (BLB: 50 mM Tris-HCl pH 8.0, 300 mM NaCl, 10 mM MgCl_2_, 10 mM Imidazole, 10% glycerol, 5 mM 2-ME, 1 mM PMSF, 1× cOmplete™ protease inhibitor cocktail [Roche, 11873580001] and 1 ng/µL Pepstatin), and lysed using a High-Pressure Cell Press Homogenizer‎ (Avestin Emulsiflex-C5). The lysate was supplemented with 0.1 mg/ml DNase I, 0.1 mg/mL RNase A, and 0.1% Triton-X 100 and cleared by centrifugation at 30,000g for 1 hr at 4°C. The supernatant was applied to an NTA-Ni column (Qiagen, 30230), and washed three times with BLB supplied with gradually ascent concentration of imidazole (10 mM, 25 mM, 50 mM) and gradually descent concentration of Sodium Chloride (500 mM, 300 mM, 150 mM). The proteins with 6×His were eluted with 50 mM Tris-HCl pH 8.0, 150 mM NaCl, 250 mM Imidazole, 10% glycerol and 5 mM 2-ME. The eluted proteins were separated by Superdex 200 Increase 10/300 GL column (GE Healthcare) equilibrated in size exclusive buffer (SEC: 50 mM Tris-HCl pH 7.5, 150 mM NaCl, 10% glycerol, 2mM 2-ME). Finally, purified proteins were concentrated in SEC and frozen in liquid nitrogen. The concentration of the proteins were determined by Bradford Assay ^7^, and the purity of the proteins were qualified by SDS-PAGE and Coomassie Bright Blue R-250 staining (Fig. S3C).

**IP of proteins expressed in yeast**

To isolate the Rim4-FLAG complex, frozen cell lysate powder (~ 500 OD_600_ units) was thawed in 2ml 1 × IP buffer (50 mM HEPES-KOH (pH 6.8), 150 mM KOAc, 2 mM Mg(OAc)_2_, 1 mM CaCl2, 15% glycerol, 1% NP-40, 1× cOmplete™ protease inhibitor cocktail [Roche, 11873580001], 1× PhosStop™ phosphatase inhibitors[Roche, 4906845001]) and cleared twice by centrifugation at 1,000 × g for 5 min at 4°C. 10 µl of Protein G Dynabeads (Invitrogen) that were loaded with mouse anti-FLAG M2 antibody (Sigma, F3165) were then added to the cell extract, incubated for 3 h at 4°C with constant agitation. The beads were collected, washed 5 times with IP buffer, and bound proteins were eluted with 25 µl 1 mg/ml 3 × FLAG peptide (Sigma, F4799) at 4°C. Eluates were aliquoted, frozen in liquid nitrogen, and stored at -80°C. The purified Rim4-FLAG complex was resolved by SDS-PAGE and analyzed by SYPRO Ruby staining (Thermo Fisher, S12000).

**Co-IP and Pull-Down Assays**

IP of proteins expressed in yeast Cells grown in SPM or YPD medium were pelleted at 3,000 × g for 5 min, 4°C, washed with ice-cold distilled water containing 1mM PMSF. The pellets were resuspended in ice-cold yeast lysis buffer (YLB: 50mM HEPES-KOH, pH 6.8, 150 mM KOAc, 2 mM MgCl_2_, 1 mM CaCl_2_, 0.2 M sorbitol, 10mM PMSF, 2× protease inhibitor cocktails [Roche, 11873580001]), dropped into liquid nitrogen, ground using a Retsch ball mill (PM100 or MM400) and stored at -80°C for immunoprecipitation (IP), as well as for atg1 kinase assay and immunoblotting (IB).

**Immunoblotting**

1.4 OD_600_ of cells in each condition were collected by pelleting at 3,000 × g for 5 min, RT, and incubated in water containing 10 mM PMSF at RT for 5 min. 2× SDS Loading Buffer (125 mM Tris-HCl, pH 6.8, 4% SDS, 0.1% BPB, 20% Glycerol, 10% 2-ME) was added to the cell pellets to the final concentration of 1×. Heated at 70˚C for 5 min. The supernatants were separated by SDS-PAGE (30 min at 200 V) using the 4-20% gradient PAGE gels (26-well: Criterion™ Tris-Glycine [TGX] Stain-Free™ gels [Bio-Rad, 5678095]; 15-well: SuperPAGE™ Bis-Tris gels [GenScript, M00657]). Subsequentially, the protein samples in the gels were electroblotted onto nitrocellulose membranes (Bio-Rad, 1620115) using a Trans-Blot® semi-dry transferring cell (Bio-Rad, 1703940).

Transferred membranes were stained with 0.1% Ponceau S solution. After that, block the membrane in 5% skim milk or 1% BSA in TBST (20 mM Tris-HCl, pH 7.5, 150 mM NaCl, 0.5% Tween-20). Next, incubate sequentially in primary antibodies and HRP-, StarBright B700- or Alexa Fluor 488-conjugated secondary antibodies. Ponceau S staining and IB images were captured by ChemiDoc™ MP imaging system (Bio-Rad, 12003154).

**Endogenous Rim4 concentration determine**

1.4 OD_600_ yeast cells harboring Rim4 with C-terminal tagged EGFP (Rim4-EGFP) in SPM were collected at 0.5 hr, 10 hr, 12 hr, 14 hr and 16 hr, treat with PMSF as described above.

Then, we expressed and purified recombinant Rim4-EGFP-6×His with the same linker used in the genomic version of Rim4-EGFP between the Rim4 EGFP. Dilute the protein into 500 nM, and do a serial half dilution for 9 times, until the last concentration is 500/2^9^ nM (0.9765625 nM). Discard a half volume of the last dilution to maintain the volumes in the 10 dilution gradients were the same.

Next, add 100 µL 1× SDS loading buffer to the cell samples. And supply the recombinant protein samples with identical volume of the proteins of 2× SDS Loading buffer. Load 10 µL of both cell and protein samples on the same PAGE gel. Thus, in each well, there were cell samples equivalent to 0.14 OD_600_; and 2.5, 1.25, 0.625, 0.3125…0.00488 pmol recombinant protein in each well. Blot to the same NC membrane after gel running. Detect the endogenous Rim4-EGFP and recombinant Rim4-GFP-6×His on the membrane by Western Blot.

Quantify the bands according to the Quantification and Statistics section below. Plot the band volumes vs amount (pmol) of the recombinant proteins in each lane on X-Y coordinate. Do a simple linear regression, and the fit equation is:

$$y=1.846\times{10}^{8}x-1.387\times{10}^{5}, R^{2}=0.9985$$

Here, y presents the band volumes and x presents the amount (pmol) of the recombinant proteins in each lane. Do a transpose for later conversion of band volume to amount:

$$y=5.418\times{10}^{-9}x+9.953\times{10}^{-4}$$

Here, y presents the amount (pmol) of the recombinant proteins in each lane and x presents the band volumes.

Based on the equation and the band volumes of the endogenous Rim4-GFP bands, calculate the concentration range of Rim4-GFP in each lane was: 0.05-0.41 pmol.

It was reported that the average volume of diploid yeast cells was 8.2×10^-8^ µL ^8^ while the cell density of 1 OD_600_ yeast cell was 3×10^7^ cell/mL ^9^. Therefore, the volume of yeast cell content in each lane (10 µL sample) was 1.4 OD_600_ × 3×10^7^ cell/mL/OD_600_ × 8.2×10^-8^ µL/cell × 1000 µL/mL = 0.34 µL.

Hence, based on our calculation and knowledge, we finally calculated the average concentration of endogenous Rim4-GFP in one cell ranged from 0.23 to 1.21 µM.

**Fluorescence Microscopy Imaging**

200 µL cells growing in SPM were concentrated to 10 µL. Drop 5 µL to the glass slides (Corning, 2975-246). The cells were then covered by a piece of agar containing the same media and immediately observed under Spinning disk Confocal (Yokogawa Spinning Disk Confocal CSU-W1) Zeiss Axio Observer microscope supplied with Hamamatsu Orca-Fusion sCMOS camera and a Zeiss Plan Apochromat 63×/0.9-NA oil-immersion objective. Exposure time was 200 ms for both red (561-617/71 nm) and green (473-525/50 nm) channel (laser power 100%). Images were captured with SlideBook 6 software. Z-stack of 5 planes, 1 µm/plane, were taken for each field (202.44 µm×202.44 µm), each channel. 3 fields containing more than 300 cells were taken for each sample. The images were split for the best focus, colored, analyzed by ImageJ (version 1.53u).

**Mass Spectrometry**

The cells harboring *pZEV:Rim4-FLAG* were driven to sporulation, β-estradiol, the inducer of ZEV promoter was added at 0 hr (immediate after cells transferred into SPM). Cells were harvested at 12 hr and snap frozen to liquid nitrogen. Smash the cells by ball milling. Next, the cell lysate powder was dissolved in 1× IP buffer (50 mM HEPES-KOH, pH 6.8, 150 mM KOAc, 2 mM Mg(OAc)_2_, 1 mM CaCl_2_, 15% glycerol, 1% NP-40, 1× cOmplete™ protease inhibitor cocktail [Roche, 11873580001], 1× PhosStop™ phosphatase inhibitors[Roche, 4906845001]), followed by centrifugation at 100,000 × g for 49 min, 4°C, to separate the cytosolic fraction (supernatant, S_100_) and membrane fraction (pellet, including nuclear, organelles and membranes, P_100_). The cytosolic fraction (S_100_) and detergent dissolved membrane fraction (P_100_) were incubated with the anti-FLAG M2 monoclonal antibody bound Protein G resin on ice for 30 min, respectively. The beads were then collected and washed 5 times with IP buffer. The eluents by 25 µL 1 mg/mL 3×FLAG peptides were sent to UT Southwestern Medical Center Proteomics Core Facility for mass spectrometry analysis.

**Mass Photometry**

The cytosolic fraction of the samples sent for mass spectrometry was sent to the UT Southwestern Medical Center Macromolecular Biophysics Resource for mass photometry analysis. The result of the counts of particles were plotted on histogram by the molecular mass. The peaks were Gaussian fitted, and the mean molecular mass and standard deviation (σ) were calculated.

***in vitro* Phosphorylation and Dephosphorylation of recombinant proteins**

To conduct the *in vitro* phosphorylation of Rim4, 6 µM recombinant Rim4-6×His was incubated with 0.5 μg/μL PKA catalytic subunit (Sigma, P2645) or 0.6 µM Ime2st (∆C241) (purified in Wang Lab) in reaction buffer (50 mM HEPES-NaOH, pH 6.8, 150 mM KOAc, 2 mM Mg[OAc]_2_, 1 mM CaCl_2_, 1% NP-40, 5 mM NaF, 50 mM β-Glycerophosphate, 10 mM Na_3_VO_4_, 50 mM Na_2_PPi [Disodium pyrophosphate], 1 ng/μL Pepstatin, 1× cOmplete Protease Inhibitor Cocktail [Roche, 11873580001], 1 mM PMSF, 15% Glycerol) at RT for 1 hr.

Rim4-p in the cell lysate were first immobilized on beads, either by pulling down via Bmh1/2-GFP-6His immobilized on Ni-NTA resin (EGFP-Rim4) or by directly immobilizing on anti-FLAG M2 monoclonal antibody bound Protein G dynabeads (Rim4-3FLAG). Then the Rim4-p proteins on beads were de-phosphorylated by incubating with 1.5 µM 3FLAG-Cdc14 in reaction buffer (50 mM HEPES-NaOH, pH 6.8, 150 mM KOAc, 2 mM Mg[OAc]_2_, 1 mM CaCl_2_, 0.075% NP-40, 1 mM DTT, 1 ng/μL Pepstatin, 1× cOmplete Protease Inhibitor Cocktail [Roche, 11697498001], 1 mM PMSF, 15% Glycerol) RT for 30 min.

**ATG1 Kinase Activity Assay**

The ATG1 kinase activity assay was described in our previous study^10^. Briefly, the cell lysate was mixed with 1 × kinase buffer (150 mM KOAc, 10 mM Mg[OAc]_2_, 0.5 mM EGTA, 5 mM NaCl, 20 mM HEPES-KOH [pH 7.3], 5% glycerol) in equal volume (wt/vol) on ice and thawed by pipetting. After spin at 1000×g for 5 min twice, supernatants were mixed with different concentration of recombinant Rim4 and equal volumes of 2 × kinase mix (kinase buffer, energy mix [90 mM creatine phosphate, 2.2 mM ATP, 0.45 mg/ml creatine kinase] and 0.2 mM N^6^-phenylethyl-ATPγS [N^6^-PhEt-ATPγS, Axxora, BLG-P026-05]) and incubated for 1.5 h at room temperature. Reactions were quenched with 20 mM EDTA and then alkylated with 2.5 mM *p*-Nitrobenzyl Mesylate (PNBM, Abcam, ab138910) for 45 min at room temperature, heated in sample loading buffer, and analyzed by SDS-PAGE and following immunoblotting assay. Thiophosphorylated substrates were identified by immunoblotting with a rabbit anti-thiophosphate ester primary antibody [51-8] (Abcam, ab92570) and StarBright® B700 labeled goat anti-Rabbit IgG secondary antibody. Blot imaging was done using a Bio-Rad ChemiDoc™ MP Imaging System.

***in silico* predictions**

The functional domains of Rim4 were mapped by Simple Modular Architecture Research Tool (SMART, <http://smart.embl-heidelberg.de/smart/set_mode.cgi?NORMAL=1>).

The disordered regions of Rim4 were predicted by DISOPRED3 program (<http://bioinf.cs.ucl.ac.uk/psipred/>, check the DISOPRED3 analysis). The PBDATA file were downloaded and the scores of each position were extracted and plot on a Cartesian coordinate system.

The RNA binding sites in the C-terminus LCD areas were predicted by ProNA2020 program (<https://predictprotein.org/>).

The 14-3-3 binding sites were predicted by 14-3-3-pred program (<http://www.compbio.dundee.ac.uk/1433pred/>). The prediction was done with 3 methods: Artificial Neural Network (ANN, cut-off=0.55), Position-Specific Scoring Matrix (PSSM, cut-off = 0.8) and Support Vector Machine (SVM, cut-off = 0.25). The position of all S and T residues were calculated and scored. The 5 sites with all three method scores above the cut-off were selected as positive sites.

The full length Rim4 protein (entry: P38741) structure was predicted by AlphaFold (<https://www.alphafold.ebi.ac.uk/>). The structure of RRMs of Rim4 were predicted by Robetta program with the RoseTTAFold method (<https://robetta.bakerlab.org/>). All the predicted structures were downloaded as PDB files, visualized and analyzed by Chimera software (version 1.16) (<https://www.cgl.ucsf.edu/chimera/>).

**Quantification and statistics**

**Immunoblotting quantification**

The band volumes of the immunoblotting were analyzed by Image Lab software (Bio-Rad, version 6.1). All band volumes of the protein of interest were normalized by that of the internal reference protein Pgk1 or Hxk1. The EGFP-Rim4 variants autophagic degradation were calculated by free EGFP signals divided by total EGFP signals, including full length EGFP-Rim4 variants and the free EGFP signals.

**Sporulation efficiency calculation**

At 60 hr after transferring into SPM, cells were collected and diluted accordingly. Added the samples into a hemocytometer chamber (Hausser Scientific, 3110) and count the total cell numbers (>300) as well as the tetrads numbers. The sporulation efficiency is presented by the tetrad percentage, i.e., tetrads number divided by total cell number.

**FM analysis**

FM images were analyzed by ImageJ (1.53u). The Min and Max Display Value of the mScarlet channel were set as (0, 300); that of the EGFP channel were set as (0, 500) unless otherwise indicated. The layer best fit the focus were split and colored with the LUT: mScarlet-red; EGFP-green; bright filed-gray. The merged images were created with the above LUT and saved as a separate file.

1.	Blethrow, J., Zhang, C., Shokat, K.M., and Weiss, E.L. (2004). Design and use of analog-sensitive protein kinases. Curr Protoc Mol Biol *Chapter 18*, Unit 18 11. 10.1002/0471142727.mb1811s66.

7.	Kielkopf, C.L., Bauer, W., and Urbatsch, I.L. (2020). Bradford Assay for Determining Protein Concentration. Cold Spring Harb Protoc *2020*, 102269. 10.1101/pdb.prot102269.

8.	Jorgensen, P., Nishikawa, J.L., Breitkreutz, B.J., and Tyers, M. (2002). Systematic identification of pathways that couple cell growth and division in yeast. Science *297*, 395-400. 10.1126/science.1070850.

9.	Day, A., Schneider, C., and Schneider, B.L. (2004). Yeast cell synchronization. Methods Mol Biol *241*, 55-76. 10.1385/1-59259-646-0:55.

10.	Feng, W., Argüello-Miranda, O., Qian, S., and Wang, F. (2022). Cdc14 spatiotemporally dephosphorylates Atg13 to activate autophagy during meiotic divisions. Journal of Cell Biology *221*. 10.1083/jcb.202107151.
