## Supplementary material for "Cdc14 spatiotemporally regulates Rim4-mRNA complex assembly and stability during meiosis": Key Resources Table

| REAGENT or RESOURCE | SOURCE | IDENTIFIER |
| --- | --- | --- |
| Antibodies | | |
| Monoclonal anti-GFP mouse IgG (1:10,000) | Roche | Cat#11814460001; RRID: AB_390913 |
| Monoclonal anti-FLAG mouse IgG (1:5,000) | Sigma | Cat# F3165;  RRID: AB_259529 |
| Monoclonal anti-Phospho-PKA Substrate (RRxS*/T*) Rabbit IgG (1:1,000) | Cell Signaling Technologies | Cat# 9624S;  RRID: AB_331817 |
| Recombinant anti-Thiophosphate ester Rabbit IgG (1:3,000) | Abcam | Cat# ab92570;  RRID: AB_10562142 |
| Polyclonal anti-hexokinase 1 Rabbit IgG (1:10,000) | United States Biological | Cat# 169073 |
| Polyclonal anti-PGK1 Rabbit IgG (1:10,000) | Antibodies Online Inc. | Cat# ABIN568371  RRID: AB_2924789 |
| StarBright® B700 labeled goat anti-mouse IgG secondary antibody (1:10,000) | Bio-Rad | Cat# 12004158  RRID: AB_2884948 |
| Alexa Fluor® 488 labeled goat anti-rabbit IgG secondary (1:10,000) | Thermo Fisher Scientific | Cat# A11034;  RRID: AB_2576217 |
| ECL-anti-rabbit HRP IgG (1:10,000) | Cytiva | Cat# NA9340;  RRID: AB_772191 |
| Chemicals, Peptides, and Recombinant Proteins | | |
| Bacto Peptone | BD Biosciences | Cat# 211820 |
| Bacto Yeast Extract | BD Biosciences | Cat# 212720 |
| Glucose/Dextrose | Fisher BioReagents | Cat# D16-10 |
| Potassium Acetate | Fisher BioReagents | Cat# P171-500 |
| Yeast Nitro Base w/o amino acid | VWR | Cat# 90004-146 |
| Bacto Agar | VWR | Cat# 90000-762 |
| β-estradiol | Sigma | Cat# E8875 |
| 1NM-PP1 | APExBIO | Cat# B1299 |
| Glycerol | Sigma | Cat# 356352 |
| IPTG (dioxane free) | US Biological | Cat# I8500 |
| G418 disulfate salt | Sigma | Cat# A1720 |
| Nourseothricin Sulfate | Gold Biotechnology | Cat# N-500-1 |
| Ni-NTA agarose | Qiagen | Cat# 30230 |
| Sodium Citrate | Fisher Scientific | Cat# BP327-500 |
| Tris Base | Sigma | Cat# T1378 |
| Sodium Chloride | Sigma | Cat# S7653 |
| Magnesium Chloride | Sigma | Cat# M2670 |
| 2-Mercaptoethanol | Sigma | Cat# 63689 |
| Imidazole | Fisher Scientific | Cat# ICN1020335 |
| Dynabeads Protein G | Fisher Scientific | Cat# 10004D |
| Pepstatin | Roche | Cat# 11359053001 |
| Protease Inhibitor Cocktail | Roche | Cat# 11873580001 |
| PMSF | Sigma | Cat# 78830 |
| Sodium Dodecyl Sulfate | VWR | Cat# EM-7910 |
| Glycine | Sigma | Cat# G7126 |
| Tween-20 | Sigma | Cat# P1379 |
| Bromophenol Blue | Sigma | Cat# B0126 |
| Skim milk powder | VWR | Cat# IC90288705 |
| Bovine Serum Albumin | Sigma | Cat# A2153 |
| Sodium Azide | Sigma | Cat# S2002 |
| 4-20% Criterion™ TGX Stain-Free™ Gel | Bio-Rad | Cat# 5678095 |
| SurePAGE™ 4-20% Bis-Tris Gel | Genscript | Cat# M00657 |
| 3×FLAG peptides | Sigma | Cat# F4799 |
| λ-Phosphatase | NEB | Cat# P0753S |
| Zymolyse (20T) | AMS Bio | Cat# 120491-1 |
| Protein Kinase A Catalytic Subunit from bovine heart | Sigma | Cat# P2645 |
| Rim4-6His | This study | NA |
| Rim4-GFP-6His | This study | NA |
| Rim4(47A)-6His | This study | NA |
| Rim4(PxLm)-6His | This study | NA |
| Bmh1-GFP-6His | This study | NA |
| Bmh2-GFP-6His | This study | NA |
| Bmh1-3FLAG-6His | This study | NA |
| 6His-3FLAG-Cdc14 | ^1^ | NA |
| Experimental Models: Organisms/Strains | | |
| See table S1 |  |  |
| Recombinant DNA | | |
| See table S2 |  |  |
| Software and Algorithms | | |
| Image Lab™ (version 6.1) | Bio-Rad | https://www.bio-rad.com/en-us/product/image-lab-software |
| Image J (version 1.53u) | National Institutes of Health (Public Domain) | <https://imagej-nih-gov.foyer.swmed.edu/ij/> |
| SlideBook 6 | Intelligent imaging | https://www.intelligent-imaging.com/slidebook |
| SnapGene (version 6.1.2) | Dotmatic | https://www.snapgene.com/ |
| GraphPad Prism (version 9.4.1) | Dotmatic | https://www.graphpad.com/ |
| AlphaFold | DeepMind Technologies | https://alphafold.com/ |
| Robetta | Baker Lab, HHMI’s Janelia Research Campus | http://www.robetta.org/ |
| DISOPRED 3.16 | UCL Department of Computer Science | http://bioinf.cs.ucl.ac.uk/psipred/ |

1.	Feng, W., Argüello-Miranda, O., Qian, S., and Wang, F. (2022). Cdc14 spatiotemporally dephosphorylates Atg13 to activate autophagy during meiotic divisions. Journal of Cell Biology *221*. 10.1083/jcb.202107151.
