## Supplementary material for "Cdc14 spatiotemporally regulates Rim4-mRNA complex assembly and stability during meiosis": Table S1

Table S1. Strains used in this study. Strains are derivatives of W303 (*ade2-1 his3-11,15 leu2-3,112 trp1-1 ura3-1 can1-100*)

| Strain Name | Figures | Genotype | Source |
| --- | --- | --- | --- |
| FWY2331 | 1A, 1B, 1C, S1B, 3A, 4F, 5B, 5C, 5D, 5E, 6C, 6D | MAT A/α, Atg1(M_102_G)/Atg1(M_102_G), Ndt80::P_GAL1_-Ndt80:KanMX/Ndt80::P_GAL1_-Ndt80:KanMX, leu2::P_ACT1_-Gal4(848).ER:NatMX/leu2::P_ACT1_-Gal4(848).ER:NatMX, ∆rim4::Ura3/∆rim4::Ura3, his3::P_RIM4_-EGFP-Rim4:His3/his3, trp1::P_NUP49_-Nup49-mScarlet:Trp1/trp1 | This study |
| FWY2373 | 1A, 1B, S1B | MAT A/α, Atg1(M_102_G)/Atg1(M_102_G), Ndt80::P_GAL1_-Ndt80:KanMX/Ndt80::P_GAL1_-Ndt80:KanMX, leu2::P_ACT1_-Gal4(848).ER:NatMX/leu2::P_ACT1_-Gal4(848).ER:NatMX, ∆rim4::Ura3/∆rim4::Ura3, his3::P_RIM4_-EGFP-Rim4(R1A):His3/his3, trp1::P_NUP49_-Nup49-mScarlet:Trp1/trp1 | This study |
| FWY2374 | 1A, 1B, 1C, S1B | MAT A/α, Atg1(M_102_G)/Atg1(M_102_G), Ndt80::P_GAL1_-Ndt80:KanMX/Ndt80::P_GAL1_-Ndt80:KanMX, leu2::P_ACT1_-Gal4(848).ER:NatMX/leu2::P_ACT1_-Gal4(848).ER:NatMX, ∆rim4::Ura3/∆rim4::Ura3, his3::P_RIM4_-EGFP-Rim4(R2A):His3/his3, trp1::P_NUP49_-Nup49-mScarlet:Trp1/trp1 | This study |
| FWY2375 | 1A, 1B, 1C, S1B | MAT A/α, Atg1(M_102_G)/Atg1(M_102_G), Ndt80::P_GAL1_-Ndt80:KanMX/Ndt80::P_GAL1_-Ndt80:KanMX, leu2::P_ACT1_-Gal4(848).ER:NatMX/leu2::P_ACT1_-Gal4(848).ER:NatMX, ∆rim4::Ura3/∆rim4::Ura3, his3::P_RIM4_-EGFP-Rim4(R3A):His3/his3, trp1::P_NUP49_-Nup49-mScarlet:Trp1/trp1 | This study |
| FWY2376 | 1A, 1B, S1B | MAT A/α, Atg1(M_102_G)/Atg1(M_102_G), Ndt80::P_GAL1_-Ndt80:KanMX/Ndt80::P_GAL1_-Ndt80:KanMX, leu2::P_ACT1_-Gal4(848).ER:NatMX/leu2::P_ACT1_-Gal4(848).ER:NatMX, ∆rim4::Ura3/∆rim4::Ura3, his3::P_RIM4_-EGFP-Rim4(R4A):His3/his3, trp1::P_NUP49_-Nup49-mScarlet:Trp1/trp1 | This study |
| FWY2282 | 1A, 1B, 1C, S1B | MAT A/α, Atg1(M_102_G)/Atg1(M_102_G), Ndt80::P_GAL1_-Ndt80:KanMX/Ndt80::P_GAL1_-Ndt80:KanMX, leu2::P_ACT1_-Gal4(848).ER:NatMX/leu2::P_ACT1_-Gal4(848).ER:NatMX, ∆rim4::Ura3/∆rim4::Ura3, his3::P_RIM4_-EGFP-Rim4(R5A):His3/his3, trp1::P_NUP49_-Nup49-mScarlet:Trp1/trp1 | This study |
| FWY2377 | 1A, 1B, S1B | MAT A/α, Atg1(M_102_G)/Atg1(M_102_G), Ndt80::P_GAL1_-Ndt80:KanMX/Ndt80::P_GAL1_-Ndt80:KanMX, leu2::P_ACT1_-Gal4(848).ER:NatMX/leu2::P_ACT1_-Gal4(848).ER:NatMX, ∆rim4::Ura3/∆rim4::Ura3, his3::P_RIM4_-EGFP-Rim4(R6A):His3/his3, trp1::P_NUP49_-Nup49-mScarlet:Trp1/trp1 | This study |
| FWY2378 | 1A, 1B, S1B | MAT A/α, Atg1(M_102_G)/Atg1(M_102_G), Ndt80::P_GAL1_-Ndt80:KanMX/Ndt80::P_GAL1_-Ndt80:KanMX, leu2::P_ACT1_-Gal4(848).ER:NatMX/leu2::P_ACT1_-Gal4(848).ER:NatMX, ∆rim4::Ura3/∆rim4::Ura3, his3::P_RIM4_-EGFP-Rim4(R7A):His3/his3, trp1::P_NUP49_-Nup49-mScarlet:Trp1/trp1 | This study |
| FWY2379 | 1A, 1B, S1B | MAT A/α, Atg1(M_102_G)/Atg1(M_102_G), Ndt80::P_GAL1_-Ndt80:KanMX/Ndt80::P_GAL1_-Ndt80:KanMX, leu2::P_ACT1_-Gal4(848).ER:NatMX/leu2::P_ACT1_-Gal4(848).ER:NatMX, ∆rim4::Ura3/∆rim4::Ura3, his3::P_RIM4_-EGFP-Rim4(R8A):His3/his3, trp1::P_NUP49_-Nup49-mScarlet:Trp1/trp1 | This study |
| FWY2380 | 1A, 1B, S1B | MAT A/α, Atg1(M_102_G)/Atg1(M_102_G), Ndt80::P_GAL1_-Ndt80:KanMX/Ndt80::P_GAL1_-Ndt80:KanMX, leu2::P_ACT1_-Gal4(848).ER:NatMX/leu2::P_ACT1_-Gal4(848).ER:NatMX, ∆rim4::Ura3/∆rim4::Ura3, his3::P_RIM4_-EGFP-Rim4(R9A):His3/his3, trp1::P_NUP49_-Nup49-mScarlet:Trp1/trp1 | This study |
| FWY2365 | 1A, 1B, S1B | MAT A/α, Atg1(M_102_G)/Atg1(M_102_G), Ndt80::P_GAL1_-Ndt80:KanMX/Ndt80::P_GAL1_-Ndt80:KanMX, leu2::P_ACT1_-Gal4(848).ER:NatMX/leu2::P_ACT1_-Gal4(848).ER:NatMX, ∆rim4::Ura3/∆rim4::Ura3, his3::P_RIM4_-EGFP-Rim4(R1E):His3/his3, trp1::P_NUP49_-Nup49-mScarlet:Trp1/trp1 | This study |
| FWY2366 | 1A, 1B, S1B | MAT A/α, Atg1(M_102_G)/Atg1(M_102_G), Ndt80::P_GAL1_-Ndt80:KanMX/Ndt80::P_GAL1_-Ndt80:KanMX, leu2::P_ACT1_-Gal4(848).ER:NatMX/leu2::P_ACT1_-Gal4(848).ER:NatMX, ∆rim4::Ura3/∆rim4::Ura3, his3::P_RIM4_-EGFP-Rim4(R2E):His3/his3, trp1::P_NUP49_-Nup49-mScarlet:Trp1/trp1 | This study |
| FWY2367 | 1A, 1B, S1B | MAT A/α, Atg1(M_102_G)/Atg1(M_102_G), Ndt80::P_GAL1_-Ndt80:KanMX/Ndt80::P_GAL1_-Ndt80:KanMX, leu2::P_ACT1_-Gal4(848).ER:NatMX/leu2::P_ACT1_-Gal4(848).ER:NatMX, ∆rim4::Ura3/∆rim4::Ura3, his3::P_RIM4_-EGFP-Rim4(R3E):His3/his3, trp1::P_NUP49_-Nup49-mScarlet:Trp1/trp1 | This study |
| FWY2368 | 1A, 1B, S1B | MAT A/α, Atg1(M_102_G)/Atg1(M_102_G), Ndt80::P_GAL1_-Ndt80:KanMX/Ndt80::P_GAL1_-Ndt80:KanMX, leu2::P_ACT1_-Gal4(848).ER:NatMX/leu2::P_ACT1_-Gal4(848).ER:NatMX, ∆rim4::Ura3/∆rim4::Ura3, his3::P_RIM4_-EGFP-Rim4(R4E):His3/ his3, trp1::P_NUP49_-Nup49-mScarlet:Trp1/trp1 | This study |
| FWY2326 | 1A, 1B, 1E, S1B | MAT A/α, Atg1(M_102_G)/Atg1(M_102_G), Ndt80::P_GAL1_-Ndt80:KanMX/Ndt80::P_GAL1_-Ndt80:KanMX, leu2::P_ACT1_-Gal4(848).ER:NatMX/leu2::P_ACT1_-Gal4(848).ER:NatMX, ∆rim4::Ura3/∆rim4::Ura3, his3::P_RIM4_-EGFP-Rim4(R5E):His3/his3::P_RIM4_-EGFP-Rim4(R5E):His3, trp1::P_NUP49_-Nup49-mScarlet:Trp1/trp1 | This study |
| FWY2369 | 1A, 1B, 1E, S1B | MAT A/α, Atg1(M_102_G)/Atg1(M_102_G), Ndt80::P_GAL1_-Ndt80:KanMX/Ndt80::P_GAL1_-Ndt80:KanMX, leu2::P_ACT1_-Gal4(848).ER:NatMX/leu2::P_ACT1_-Gal4(848).ER:NatMX, ∆rim4::Ura3/∆rim4::Ura3, his3::P_RIM4_-EGFP-Rim4(R6E):His3/his3, trp1::P_NUP49_-Nup49-mScarlet:Trp1/trp1 | This study |
| FWY2370 | 1A, 1B, 1E, S1B | MAT A/α, Atg1(M_102_G)/Atg1(M_102_G), Ndt80::P_GAL1_-Ndt80:KanMX/Ndt80::P_GAL1_-Ndt80:KanMX, leu2::P_ACT1_-Gal4(848).ER:NatMX/leu2::P_ACT1_-Gal4(848).ER:NatMX, ∆rim4::Ura3/∆rim4::Ura3, his3::P_RIM4_-EGFP-Rim4(R7E):His3/his3, trp1::P_NUP49_-Nup49-mScarlet:Trp1/trp1 | This study |
| FWY2371 | 1A, 1B, S1B | MAT A/α, Atg1(M_102_G)/Atg1(M_102_G), Ndt80::P_GAL1_-Ndt80:KanMX/Ndt80::P_GAL1_-Ndt80:KanMX, leu2::P_ACT1_-Gal4(848).ER:NatMX/leu2::P_ACT1_-Gal4(848).ER:NatMX, ∆rim4::Ura3/∆rim4::Ura3, his3::P_RIM4_-EGFP-Rim4(R8E):His3/his3, trp1::P_NUP49_-Nup49-mScarlet:Trp1/trp1 | This study |
| FWY2372 | 1A, 1B, S1B | MAT A/α, Atg1(M_102_G)/Atg1(M_102_G), Ndt80::P_GAL1_-Ndt80:KanMX/Ndt80::P_GAL1_-Ndt80:KanMX, leu2::P_ACT1_-Gal4(848).ER:NatMX/leu2::P_ACT1_-Gal4(848).ER:NatMX, ∆rim4::Ura3/∆rim4::Ura3, his3::P_RIM4_-EGFP-Rim4(R9E):His3/his3, trp1::P_NUP49_-Nup49-mScarlet:Trp1/trp1 | This study |
| FWY1170 | 2A, 2B, 2C, S5A | MAT A/α, Atg1(M_102_G)/Atg1(M_102_G), ∆rim4::Ura3/∆rim4::Ura3, leu2::P_ZEV_-Rim4-3×FLAG:Leu2/leu2::P_ZEV_-Rim4-3×FLAG:Leu2 | This study |
| FWY1471 | 2E, 2F, 2G, 3D, 4A, 4D, 4E, S4A, 5A, 5F, S5C, 6E | MAT A/α, Atg1(M_102_G)/Atg1(M_102_G), Ndt80::P_GAL1_-Ndt80:KanMX/Ndt80::P_GAL1_-Ndt80:KanMX, leu2::P_ACT1_-Gal4(848).ER:NatMX/leu2::P_ACT1_-Gal4(848).ER:NatMX, ∆rim4::Ura3/∆rim4::Ura3, his3::P_RIM4_-EGFP-Rim4:His3/his3::P_RIM4_-EGFP-Rim4:His3 | This study |
| FWY1861 | 2E, 2F | MAT A/α, Atg1(M_102_G)/Atg1(M_102_G), Ndt80::P_GAL1_-Ndt80:KanMX/Ndt80::P_GAL1_-Ndt80:KanMX, leu2::P_ACT1_-Gal4(848).ER:NatMX/leu2::P_ACT1_-Gal4(848).ER:NatMX, ∆rim4::Ura3/∆rim4::Ura3, his3::P_RIM4_-EGFP-Rim4(S_367_CT_368_C):His3/his3::P_RIM4_-EGFP-Rim4(S_367_CT_368_C):His3 | This study |
| FWY1477 | 2E, 2G, 3D | MAT A/α, Atg1(M_102_G)/Atg1(M_102_G), Ndt80::P_GAL1_-Ndt80:KanMX/Ndt80::P_GAL1_-Ndt80:KanMX, leu2::P_ACT1_-Gal4(848).ER:NatMX/leu2::P_ACT1_-Gal4(848).ER:NatMX, ∆rim4::Ura3/∆rim4::Ura3, his3::P_RIM4_-EGFP-Rim4(5C):His3/his3::P_RIM4_-EGFP-Rim4(5C):His3 | This study |
| FWY1860 | 2E | MAT A/α, Atg1(M_102_G)/Atg1(M_102_G), Ndt80::P_GAL1_-Ndt80:KanMX/Ndt80::P_GAL1_-Ndt80:KanMX, leu2::P_ACT1_-Gal4(848).ER:NatMX/leu2::P_ACT1_-Gal4(848).ER:NatMX, ∆rim4::Ura3/∆rim4::Ura3, his3::P_RIM4_-EGFP-Rim4(S_525_CS_607_C):His3/his3::P_RIM4_-EGFP-Rim4(S_525_CS_607_C):His3 | This study |
| FWY1475 | 2E, 2G | MAT A/α, Atg1(M_102_G)/Atg1(M_102_G), Ndt80::P_GAL1_-Ndt80:KanMX/Ndt80::P_GAL1_-Ndt80:KanMX, leu2::P_ACT1_-Gal4(848).ER:NatMX/leu2::P_ACT1_-Gal4(848).ER:NatMX, ∆rim4::Ura3/∆rim4::Ura3, his3::P_RIM4_-EGFP-Rim4(S_525_C):His3/his3::P_RIM4_-EGFP-Rim4(S_525_C):His3 | This study |
| FWY1863 | 2E | MAT A/α, Atg1(M_102_G)/Atg1(M_102_G), Ndt80::P_GAL1_-Ndt80:KanMX/Ndt80::P_GAL1_-Ndt80:KanMX, leu2::P_ACT1_-Gal4(848).ER:NatMX/leu2::P_ACT1_-Gal4(848).ER:NatMX, ∆rim4::Ura3/∆rim4::Ura3, his3::P_RIM4_-EGFP-Rim4(S_525_CT_216_C):His3/his3::P_RIM4_-EGFP-Rim4(S_525_CT_216_C):His3 | This study |
| FWY2226 | 2F | MAT A/α, Atg1(M_102_G)/Atg1(M_102_G), Ndt80::P_GAL1_-Ndt80:KanMX/Ndt80::P_GAL1_-Ndt80:KanMX, leu2::P_ACT1_-Gal4(848).ER:NatMX/leu2::P_ACT1_-Gal4(848).ER:NatMX, ∆rim4::Ura3/∆rim4::Ura3, his3::P_RIM4_-EGFP-Rim4(3C):His3/his3::P_RIM4_-EGFP-Rim4(3C):His3 | This study |
| FWY1472 | 2G | MAT A/α, Atg1(M_102_G)/Atg1(M_102_G), Ndt80::P_GAL1_-Ndt80:KanMX/Ndt80::P_GAL1_-Ndt80:KanMX, leu2::P_ACT1_-Gal4(848).ER:NatMX/leu2::P_ACT1_-Gal4(848).ER:NatMX, ∆rim4::Ura3/∆rim4::Ura3, his3::P_RIM4_-EGFP-Rim4(T_216_C):His3/his3::P_RIM4_-EGFP-Rim4(T_216_C):His3 | This study |
| FWY1473 | 2G | MAT A/α, Atg1(M_102_G)/Atg1(M_102_G), Ndt80::P_GAL1_-Ndt80:KanMX/Ndt80::P_GAL1_-Ndt80:KanMX, leu2::P_ACT1_-Gal4(848).ER:NatMX/leu2::P_ACT1_-Gal4(848).ER:NatMX, ∆rim4::Ura3/∆rim4::Ura3, his3::P_RIM4_-EGFP-Rim4(S_367_C):His3/his3::P_RIM4_-EGFP-Rim4(S_367_C):His3 | This study |
| FWY1474 | 2G | MAT A/α, Atg1(M_102_G)/Atg1(M_102_G), Ndt80::P_GAL1_-Ndt80:KanMX/Ndt80::P_GAL1_-Ndt80:KanMX, leu2::P_ACT1_-Gal4(848).ER:NatMX/leu2::P_ACT1_-Gal4(848).ER:NatMX, ∆rim4::Ura3/∆rim4::Ura3, his3::P_RIM4_-EGFP-Rim4(T_368_C):His3/his3::P_RIM4_-EGFP-Rim4(T_368_C):His3 | This study |
| FWY1476 | 2G | MAT A/α, Atg1(M_102_G)/Atg1(M_102_G), Ndt80::P_GAL1_-Ndt80:KanMX/Ndt80::P_GAL1_-Ndt80:KanMX, leu2::P_ACT1_-Gal4(848).ER:NatMX/leu2::P_ACT1_-Gal4(848).ER:NatMX, ∆rim4::Ura3/∆rim4::Ura3, his3::P_RIM4_-EGFP-Rim4(S_607_C):His3/his3::P_RIM4_-EGFP-Rim4(S_607_C):His3 | This study |
| FWY2339 | 3A, 5B, 6C, 6D | MAT A/α, Atg1(M_102_G)/Atg1(M_102_G), Ndt80::P_GAL1_-Ndt80:KanMX/Ndt80::P_GAL1_-Ndt80:KanMX, leu2::P_ACT1_-Gal4(848).ER:NatMX/leu2::P_ACT1_-Gal4(848).ER:NatMX, ∆rim4::Ura3/∆rim4::Ura3, his3::P_RIM4_-EGFP-Rim4(5C):His3/his3, trp1::P_NUP49_-Nup49-mScarlet:Trp1/trp1 | This study |
| FWY2280 | 4F, 5B, 5C, 5D, 6C, 6D | MAT A/α, Atg1(M_102_G)/Atg1(M_102_G), Ndt80::P_GAL1_-Ndt80:KanMX/Ndt80::P_GAL1_-Ndt80:KanMX, leu2::P_ACT1_-Gal4(848).ER:NatMX/leu2::P_ACT1_-Gal4(848).ER:NatMX, ∆rim4::Ura3/∆rim4::Ura3, his3::P_RIM4_-EGFP-Rim4(PxLm=P454AL456G):His3/his3, trp1::P_NUP49_-Nup49-mScarlet:Trp1/trp1 | This study |
| FWY1989 | 4D, 4E | MAT A/α, Atg1(M_102_G)/Atg1(M_102_G), Ndt80::P_GAL1_-Ndt80:KanMX/Ndt80::P_GAL1_-Ndt80:KanMX, leu2::P_ACT1_-Gal4(848).ER:NatMX/leu2::P_ACT1_-Gal4(848).ER:NatMX, ∆rim4::Ura3/∆rim4::Ura3, his3::P_RIM4_-EGFP-Rim4(PxL_Sic1_):His3/his3::P_RIM4_-EGFP-Rim4(PxL_Sic1_):His3 | This study |
| FWY2412 | 4F | MAT A/α, Atg1(M_102_G)/Atg1(M_102_G), Ndt80::P_GAL1_-Ndt80:KanMX/Ndt80::P_GAL1_-Ndt80:KanMX, leu2::P_ACT1_-Gal4(848).ER:NatMX/leu2::P_ACT1_-Gal4(848).ER:NatMX, ∆rim4::Ura3/∆rim4::Ura3, his3::P_RIM4_-EGFP-Rim4(PxL_Sic1_):His3/his3::P_RIM4_-EGFP-Rim4(PxL_Sic1_):His3, trp1::P_NUP49_-Nup49-mScarlet:Trp1/trp1 | This study |
| FWY1862 | 5E | MAT A/α, Atg1(M_102_G)/Atg1(M_102_G), Ndt80::P_GAL1_-Ndt80:KanMX/Ndt80::P_GAL1_-Ndt80:KanMX, leu2::P_ACT1_-Gal4(848).ER:NatMX/leu2::P_ACT1_-Gal4(848).ER:NatMX, ∆rim4::Ura3/∆rim4::Ura3, his3::P_RIM4_-EGFP-Rim4(PxLm+5C):His3/his3::P_RIM4_-EGFP-Rim4(PxLm+5C):His3 | This study |
| FWY2381 | 5B, 5C, 5D | MAT A/α, Atg1(M_102_G)/Atg1(M_102_G), Ndt80::P_GAL1_-Ndt80:KanMX/Ndt80::P_GAL1_-Ndt80:KanMX, leu2::P_ACT1_-Gal4(848).ER:NatMX/leu2::P_ACT1_-Gal4(848).ER:NatMX, ∆rim4::Ura3/∆rim4::Ura3, his3::P_RIM4_-EGFP-Rim4(PxLm+5C):His3/his3, trp1::P_NUP49_-Nup49-mScarlet:Trp1/trp1 | This study |
| FWY2334 | 1D, S5B | MAT A/α, Atg1(M_102_G)/Atg1(M_102_G), Ndt80::P_GAL1_-Ndt80:KanMX/Ndt80::P_GAL1_-Ndt80:KanMX, leu2::P_ACT1_-Gal4(848).ER:NatMX/leu2::P_ACT1_-Gal4(848).ER:NatMX, ∆rim4::Ura3/∆rim4::Ura3, his3::P_RIM4_-EGFP-Rim4(FL=F139LF349L):His3/his3, trp1::P_NUP49_-Nup49-mScarlet:Trp1/trp1 | This study |
| FWY2313 | S5B | MAT A/α, Atg1(M_102_G)/Atg1(M_102_G), Ndt80::P_GAL1_-Ndt80:KanMX/Ndt80::P_GAL1_-Ndt80:KanMX, leu2::P_ACT1_-Gal4(848).ER:NatMX/leu2::P_ACT1_-Gal4(848).ER:NatMX, ∆rim4::Ura3/∆rim4::Ura3, his3::P_RIM4_-EGFP-Rim4(FL+5C):His3/his3, trp1::P_NUP49_-Nup49-mScarlet:Trp1/trp1 | This study |
| FWY1492 | S5C | MAT A/α, Atg1(M_102_G)/Atg1(M_102_G), Ndt80::P_GAL1_-Ndt80:KanMX/Ndt80::P_GAL1_-Ndt80:KanMX, leu2::P_ACT1_-Gal4(848).ER:NatMX/leu2::P_ACT1_-Gal4(848).ER:NatMX, ∆rim4::Ura3/∆rim4::Ura3, his3::P_RIM4_-EGFP-Rim4(R6A):His3/his3::P_RIM4_-EGFP-Rim4(R6A):His3 | This study |
| FWY1864 | S5C | MAT A/α, Atg1(M_102_G)/Atg1(M_102_G), Ndt80::P_GAL1_-Ndt80:KanMX/Ndt80::P_GAL1_-Ndt80:KanMX, leu2::P_ACT1_-Gal4(848).ER:NatMX/leu2::P_ACT1_-Gal4(848).ER:NatMX, ∆rim4::Ura3/∆rim4::Ura3, his3::P_RIM4_-EGFP-Rim4(PxLm+R6A):His3/his3::P_RIM4_-EGFP-Rim4(PxLm+R6A):His3 | This study |
| FWY2353 | 5E | MAT A/α, Atg1(M_102_G)/Atg1(M_102_G), Ndt80::P_GAL1_-Ndt80:KanMX/Ndt80::P_GAL1_-Ndt80:KanMX, leu2::P_ACT1_-Gal4(848).ER:NatMX/leu2::P_ACT1_-Gal4(848).ER:NatMX, ∆rim4::Ura3/∆rim4::Ura3, his3::P_RIM4_-EGFP-Rim4(PxLm+R6A):His3/his3, trp1::P_NUP49_-Nup49-mScarlet:Trp1/trp1 | This study |
| FWY1777 | 4D, 4E, S4A, 5F, S5C, 6E | MAT A/α, Atg1(M_102_G)/Atg1(M_102_G), Ndt80::P_GAL1_-Ndt80:KanMX/Ndt80::P_GAL1_-Ndt80:KanMX, leu2::P_ACT1_-Gal4(848).ER:NatMX/leu2::P_ACT1_-Gal4(848).ER:NatMX, ∆rim4::Ura3/∆rim4::Ura3, his3::P_RIM4_-EGFP-Rim4(PxLm):His3/his3::P_RIM4_-EGFP-Rim4(PxLm):His3 | This study |
| FWY1909 | 5F, S5C | MAT A/α, Atg1(M_102_G)/Atg1(M_102_G), Ndt80::P_GAL1_-Ndt80:KanMX/Ndt80::P_GAL1_-Ndt80:KanMX, leu2::P_ACT1_-Gal4(848).ER:NatMX/leu2::P_ACT1_-Gal4(848).ER:NatMX, ∆rim4::Ura3/∆rim4::Ura3, his3::P_RIM4_-EGFP-Rim4(PxLm+SA_1_=S_532_A):His3/his3::P_RIM4_-EGFP-Rim4(PxLm+SA_1_=S_532_A):His3 | This study |
| FWY1910 | 5F, S5C | MAT A/α, Atg1(M_102_G)/Atg1(M_102_G), Ndt80::P_GAL1_-Ndt80:KanMX/Ndt80::P_GAL1_-Ndt80:KanMX, leu2::P_ACT1_-Gal4(848).ER:NatMX/leu2::P_ACT1_-Gal4(848).ER:NatMX, ∆rim4::Ura3/∆rim4::Ura3, his3::P_RIM4_-EGFP-Rim4(PxLm+SA_2_=S_658_A):His3/his3::P_RIM4_-EGFP-Rim4(PxLm+SA_2_=S_658_A):His3 | This study |
| FWY1991 | 5F, S5C | MAT A/α, Atg1(M_102_G)/Atg1(M_102_G), Ndt80::P_GAL1_-Ndt80:KanMX/Ndt80::P_GAL1_-Ndt80:KanMX, leu2::P_ACT1_-Gal4(848).ER:NatMX/leu2::P_ACT1_-Gal4(848).ER:NatMX, ∆rim4::Ura3/∆rim4::Ura3, his3::P_RIM4_-EGFP-Rim4(PxLm+SA_1/2_=S_532_AS_658_A):His3/his3::P_RIM4_-EGFP-Rim4(PxLm+SA_1/2_=S_532_AS_658_A):His3 | This study |
| FWY1990 | S5C | MAT A/α, Atg1(M_102_G)/Atg1(M_102_G), Ndt80::P_GAL1_-Ndt80:KanMX/Ndt80::P_GAL1_-Ndt80:KanMX, leu2::P_ACT1_-Gal4(848).ER:NatMX/leu2::P_ACT1_-Gal4(848).ER:NatMX, ∆rim4::Ura3/∆rim4::Ura3, his3::P_RIM4_-EGFP-Rim4(SA_1/2_=S_532_AS_658_A):His3/his3::P_RIM4_-EGFP-Rim4(SA_1/2_=S_532_AS_658_A):His3 | This study |
| FWY2413 | 6A | MAT A/α, Atg1(M_102_G)/Atg1(M_102_G), Ndt80::P_GAL1_-Ndt80:Trp1/Ndt80::P_GAL1_-Ndt80:Trp1, leu2::P_ACT1_-Gal4(848).ER:NatMX/leu2::P_ACT1_-Gal4(848).ER:NatMX, ∆rim4::Ura3/∆rim4::Ura3, his3:: P_RIM4_-mScarlet-Rim4:His3/his3:: P_RIM4_-mScarlet-Rim4:His3, trp1::P_BMH1_-Bmh1-EGFP:Trp1/trp1::P_BMH1_-Bmh1-EGFP:Trp1 | This study |
| FWY2414 | 6A | MAT A/α, Atg1(M_102_G)/Atg1(M_102_G), Ndt80::P_GAL1_-Ndt80:Trp1/Ndt80::P_GAL1_-Ndt80:Trp1, leu2::P_ACT1_-Gal4(848).ER:NatMX/leu2::P_ACT1_-Gal4(848).ER:NatMX, ∆rim4::Ura3/∆rim4::Ura3, his3:: P_RIM4_-mScarlet-Rim4:His3/his3:: P_RIM4_-mScarlet-Rim4:His3, trp1::P_BMH2_-Bmh2-EGFP:Trp1/trp1::P_BMH2_-Bmh2-EGFP:Trp1 | This study |
| FWY643 | 6B, S6C | MAT A/α, Atg1(M_102_G)/Atg1(M_102_G), Ndt80::P_GAL1_-Ndt80:Trp1/Ndt80::P_GAL1_-Ndt80:Trp1, ura3::P_TDH3_-GAL4(848).ER:Ura3/ura3::P_TDH3_-GAL4(848).ER:Ura3 | ^1^ |
| FWY486 | S6B | MAT A/α, Atg1(M_102_G)/Atg1(M_102_G), Ndt80::P_GAL1_-Ndt80:KanMX/Ndt80::P_GAL1_-Ndt80:KanMX, ura3::P_TDH3_-GAL4(848).ER:Ura3/ura3::P_TDH3_-GAL4(848).ER:Ura3, Rim4-EGFP:His3/Rim4-EGFP:His3 | This study |

1.	Feng, W., Argüello-Miranda, O., Qian, S., and Wang, F. (2022). Cdc14 spatiotemporally dephosphorylates Atg13 to activate autophagy during meiotic divisions. Journal of Cell Biology *221*. 10.1083/jcb.202107151.
