## Supplementary material for "Cdc14 spatiotemporally regulates Rim4-mRNA complex assembly and stability during meiosis": Table S2

Table S2: Plasmids used in this study.

| Plasmid Name | Description | Source |
| --- | --- | --- |
| pFW1 | pRS304>P_TDH3_-Gal4(848).ER:Ura3 | ^1^ |
| pFW127 | pAGL>P_ACT1_-Gal4(848).ER:NatMX | This study |
| pFW18 | pKL-URA>Ura3 | A gift from Denic Lab |
| pFW52 | pKT0128>yeGFP:His3 | Addgene, #8729 |
| pFW30 | P90>Rim4-allA-3V5:His3 | A gift from Berchowitz Lab |
| pFW32 | P124>Rim4-allE-3V5:His3 | A gift from Berchowitz Lab |
| pFW72 | pET29b>Rim4-6×His | This study |
| pFW95 | pET29b>Rim4-EGFP-6×His | This study |
| pFW104 | pET29b>Rim4(47A)-6×His | This study |
| pFW355 | pET29b>Rim4(5C)-6×His | This study |
| pFW81 | pET29b>Rim4(ΔC289)-6×His | This study |
| pFW354 | pET29b>Rim4(PxLm)-6×His | This study |
| pFW197 | pET29b>Bmh1-GFP-6×His | This study |
| pFW198 | pET29b>Bmh1-3×FLAG-6×His | This study |
| pFW200 | pET29b>Bmh2-GFP-6×His | This study |
| pFW394 | pET28a>6×His-3×FLAG-Cdc14 | This study |
| pFW378 | pGEX-6p-1>GST-Cdc14 | ^2^ |
| pFW45 | pNH605>P_ACT1_-Zif268-ER-VP16+P_ZEV_-Rim4-3×FLAG | This study |
| pFW338 | pRS304>P_NUP49_-Nup49-mScarlet:Trp1 | This study |
| pFW208 | pRS303>P_RIM4_-EGFP-Rim4:His3 | This study |
| pFW233 | pRS303>P_RIM4_-EGFP-Rim4(R1A):His3 | This study |
| pFW234 | pRS303>P_RIM4_-EGFP-Rim4(R2A):His3 | This study |
| pFW235 | pRS303>P_RIM4_-EGFP-Rim4(R3A):His3 | This study |
| pFW236 | pRS303>P_RIM4_-EGFP-Rim4(R4A):His3 | This study |
| pFW237 | pRS303>P_RIM4_-EGFP-Rim4(R5A):His3 | This study |
| pFW238 | pRS303>P_RIM4_-EGFP-Rim4(R6A):His3 | This study |
| pFW239 | pRS303>P_RIM4_-EGFP-Rim4(R7A):His3 | This study |
| pFW240 | pRS303>P_RIM4_-EGFP-Rim4(R8A):His3 | This study |
| pFW241 | pRS303>P_RIM4_-EGFP-Rim4(R9A):His3 | This study |
| pFW216 | pRS303>P_RIM4_-EGFP-Rim4(R1E):His3 | This study |
| pFW217 | pRS303>P_RIM4_-EGFP-Rim4(R2E):His3 | This study |
| pFW218 | pRS303>P_RIM4_-EGFP-Rim4(R3E):His3 | This study |
| pFW219 | pRS303>P_RIM4_-EGFP-Rim4(R4E):His3 | This study |
| pFW220 | pRS303>P_RIM4_-EGFP-Rim4(R5E):His3 | This study |
| pFW221 | pRS303>P_RIM4_-EGFP-Rim4(R6E):His3 | This study |
| pFW222 | pRS303>P_RIM4_-EGFP-Rim4(R7E):His3 | This study |
| pFW223 | pRS303>P_RIM4_-EGFP-Rim4(R8E):His3 | This study |
| pFW224 | pRS303>P_RIM4_-EGFP-Rim4(R9E):His3 | This study |
| pFW210 | pRS303>P_RIM4_-EGFP-Rim4(T_216_C):His3 | This study |
| pFW211 | pRS303>P_RIM4_-EGFP-Rim4(S_367_C):His3 | This study |
| pFW212 | pRS303>P_RIM4_-EGFP-Rim4(T_368_C):His3 | This study |
| pFW213 | pRS303>P_RIM4_-EGFP-Rim4(S_525_C):His3 | This study |
| pFW214 | pRS303>P_RIM4_-EGFP-Rim4(S_607_C):His3 | This study |
| pFW215 | pRS303>P_RIM4_-EGFP-Rim4(5C):His3 | This study |
| pFW285 | pRS303>P_RIM4_-EGFP-Rim4(S_525_CT_216_C):His3 | This study |
| pFW282 | pRS303>P_RIM4_-EGFP-Rim4(S_525_CS_607_C):His3 | This study |
| pFW283 | pRS303>P_RIM4_-EGFP-Rim4(S_367_CT_368_C):His3 | This study |
| pFW429 | pRS303>P_RIM4_-EGFP-Rim4(3C):His3 | This study |
| pFW250 | pRS303>P_RIM4_-EGFP-Rim4(FLm):His3 | This study |
| pFW428 | pRS303>P_RIM4_-EGFP-Rim4(FLm+5C):His3 | This study |
| pFW274 | pRS303>P_RIM4_-EGFP-Rim4(PxLm):His3 | This study |
| pFW305 | pRS303>P_RIM4_-EGFP-Rim4(PxL_Sic1_):His3 | This study |
| pFW284 | pRS303>P_RIM4_-EGFP-Rim4(PxLm+5C):His3 | This study |
| pFW286 | pRS303>P_RIM4_-EGFP-Rim4(PxLm+R6-A):His3 | This study |
| pFW294 | pRS303>P_RIM4_-EGFP-Rim4(PxLm+SA_1_):His3 | This study |
| pFW295 | pRS303>P_RIM4_-EGFP-Rim4(PxLm+SA_2_):His3 | This study |
| pFW308 | pRS303>P_RIM4_-EGFP-Rim4(SA_1/2_):His3 | This study |
| pFW311 | pRS303>P_RIM4_-EGFP-Rim4(PxLm+SA_1/2_):His3 | This study |
| pFW398 | pRS305>P_PAB1_-EGFP-Pab1:Leu2 | This study |
| pFW281 | pRS303>P_ATG8_-mScarlet-Atg8:His3 | This study |

1.	Wang, F., Zhang, R., Feng, W., Tsuchiya, D., Ballew, O., Li, J., Denic, V., and Lacefield, S. (2020). Autophagy of an Amyloid-like Translational Repressor Regulates Meiotic Exit. Dev Cell *52*, 141-151 e145. 10.1016/j.devcel.2019.12.017.

2.	Feng, W., Argüello-Miranda, O., Qian, S., and Wang, F. (2022). Cdc14 spatiotemporally dephosphorylates Atg13 to activate autophagy during meiotic divisions. Journal of Cell Biology *221*. 10.1083/jcb.202107151.
